# Structural and Boolean Network Modeling of the Levan Biosynthetic Pathway in *Bacillus subtilis*

**DOI:** 10.64898/2025.12.25.696451

**Authors:** Aruldoss Immanuel, Dharshini Priya Selvaganesan, Danthuluri Sree Lakshmi, Venkatasubramanian Ulaganathan, Ragothaman M. Yennamalli

## Abstract

**Motivation:** Levan is a fructose polymer with applications in the creation of hydrogels for drug delivery and wound healing. In industrial biotechnology. *Bacillus subtilis* is the key organism for producing levan. However, the metabolic models of *Bacillus subtilis* available do not include the biosynthesis of levan. To understand levan biosynthesis in *B. subtilis*, we employed structural systems biology integrating known structural details of proteins in the *B. subtilis* metabolic pathway to create a structure-annotated genome-scale metabolic model (GEM). To fill gaps in structural information about the enzymes, AlphaFold2 was used. Thus, this study enhances the metabolic model of *B. subtilis* by incorporating the biosynthesis of levan and including structural information about the proteins involved.

**Results:** The manually curated model links proteins and reactions to protein data bank (PDB) entries, providing structural perspectives previously overlooked in GEMs. We mapped 508 PDB structures to 168 UniProt IDs to unravel 331 out of 1250 reactions (26.5%) in *B. subtilis* with focused coverage of *sacB, sacC, sacX/Y, levD/E/F/G*, and *sacP*. The structural layer does not alter stoichiometry or constraints unless explicitly parameterized. This structure-annotated resource enables the systematic testing of phenotype predictions and design strategies. Our structure-based metabolic model advances the understanding of levan production and microbial metabolism, facilitating sustainable and efficient biotechnological processes for industrial applications.

**Availability and implementation:** Data available at the github page (https://github.com/raghuyennamalli/levan_ssbio)

## Introduction

Systems biology investigates the behavior and properties of biological systems using *in silico* modeling of molecular networks, incorporating metabolites, cofactors, and interactions. This approach helps predict the dynamics of biological pathways and can be used to modulate the production of industrially important byproducts (Breitling, 2010). Structural systems biology extends this framework by integrating molecular-level details, such as protein structures and protein-protein interactions, into genome-scale metabolic models (GEMs). This approach integrates detailed structural information with systems-level modelling to predict the behavior of complex biological networks (Brunk *et al*., 2016). Incorporating structural details into the interactome at the whole cell level provides detailed atomic-resolution insights into biological systems (Aloy and Russell, 2006).

In *Bacillus subtilis*, the research focus has been on increasing the production of value-added products such as acetoin (Renna *et al*., 1993), riboflavin and levan (Liu *et al*., 2023). Levan, a fructose polymer, is used for the formation of hydrogels, as prebiotic, as thickener, in drug delivery and in many other applications. To understand the process of levan biosynthesis, we had previously used Flux Balance Analysis (FBA) and Flux Variability Analysis (FVA) (Immanuel *et al*., 2024). In a later study, we constructed a revised enzyme-constrained genome-scale metabolic model (GEM), labeled ec_iYO844_lvn, adding levan biosynthetic reactions into the earlier model of ec_iYO844 (Massaiu *et al*., 2019). The ec_iYO844_lvn model contains 847 genes. Two genes, *pgk* and *ctaD*, were identified as potential knockout targets to increase the production of levan.

Despite the industrial importance of *B. subtilis*, the molecular networks of the bacteria remain underannotated, unlike those of other model organisms such as *Escherichia coli*. Even after the third-generation annotation of *B. subtilis*, the number of annotated genes remains relatively low. The largest available GEM (etiBsu1209) contains only 1209 genes (Bi *et al*., 2023). Moreover, most metabolic models focus on glucose as primary source and do not incorporate the reactions of the levan biosynthetic pathways. Hence, it is important to add the sucrose utilizing levan biosynthetic pathway, present in the pathway database, the Kyoto Encyclopedia of Genes and Genomes or KEGG (Kanehisa and Goto, 2000), in the metabolic modelling of *B. subtilis*.

The primary objective of this study is to improve the metabolic model of *B. subtilis* by integrating structural information with systems-level data using structural systems biology. This integration addresses the current gap in annotated information about *B. subtilis*. By enriching the *B. subtilis* model with detailed structural and gene regulatory annotations, we seek to optimize the biosynthesis of levan and other value-added products.

## 2 Methods

### Data collection and assigning structures to GEM

The UniProt database (https://www.uniprot.org/) was used as primary source for collecting the coding regions, proteome sequences, and functional annotations of *B. subtilis* strain 168 (GenBank Accession ID: GCA_000009045.1) (The UniProt Consortium, 2025). Available PDB structural data were grouped based on unique protein-coding regions, each corresponding to the unique UniProt ID based on the query *“(taxonomy_id:224308) AND (reviewed:true)”*. Each UniProt ID was subsequently matched with the corresponding *B. subtilis* gene identifier (e.g., BSU0272).

### Assigning PDB structural information to GEM

When a gene of interest is identified, we search for the protein structure encoded by that gene. Once a corresponding PDB structure is located, it is linked to gene–protein–reaction (GPR) associations. This allows us to map structural information directly to the gene. The structural data can then be integrated into the genome-scale model.

The GEM of *B. subtilis* was procured from BioModels (https://www.ebi.ac.uk/biomodels/), a database containing a repository of GEMs (Malik-Sheriff *et al*., 2020). The collected structures were manually integrated into the model, iYO844, using GPR associations as reference for each structure. PDB IDs were assigned to the reactions based on the AND rule, represented by “”. The OR rule was represented by “/”. If there were no PDB structures corresponding to the GPR gene, only the symbol was added and not the PDB ID.

### Construction of a Boolean gene regulatory network

BooleSim is a webserver designed for the simulation and analysis of Boolean networks, simplified models to represent interactions within biological systems (Bock *et al*., 2014). The rules for this network were obtained from databases for *B. subtilis*, SubtiWiki (https://subtiwiki.uni-goettingen.de/) (Pedreira *et al*., 2022), The Database of Transcriptional Regulation in *B. subtilis* (DBTBS) (https://dbtbs.hgc.jp/), and Pathway Tools (Karp *et al*., 2021). SubtiWiki is a comprehensive database of *B. subtilis*, providing curated information on its genes, proteins, pathways, and regulatory networks (Pedreira *et al*., 2022). DBTBS contains curated information on transcriptional regulation in *B. subtilis*, including data on transcription factors, promoters, regulatory interactions, and gene expression control mechanisms (Sierro *et al*., 2008).

From SubtiWiki, we obtained genes using the regulation browser tab. Sigma factor regulation and gene operon information were taken from Pathway Tools and DBTBS. The rules for the Boolean network are listed in Table 1. These rules define interactions within the network. “AND” operations are denoted by “”, “OR” operations by “||”, and “NOT” operations by “! “. Additionally, we employed “True” or ON to signify true conditions and “False” or OFF to represent false conditions. Each target node in the model is governed by a single rule expression. These rules dictate the behavior of the network, enabling us to simulate and analyze its dynamics effectively.

**Table 1.**
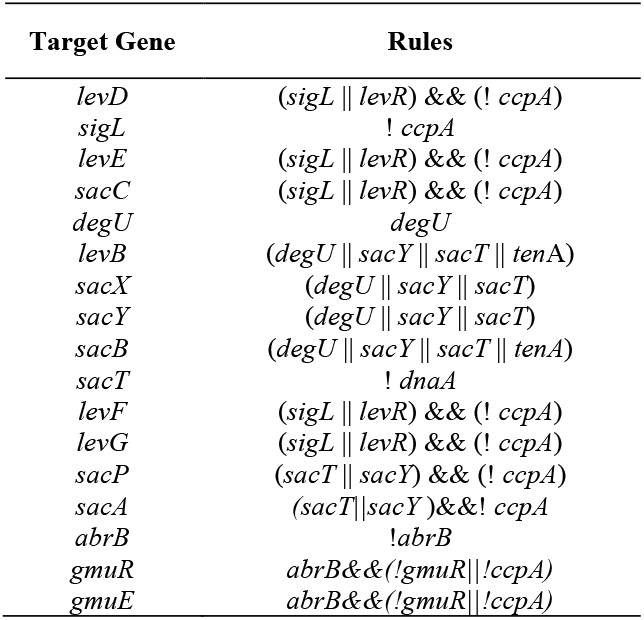
Boolean network rules for the levan biosynthetic pathway.

### Foldseek clustering to identify proteins with similar domains

Clustering analysis was carried out using the Foldseek standalone tool, the easy-cluster algorithm. This algorithm helps group protein structures with similar domains/folds into a single cluster. This facilitated the clustering of all the AlphaFold2 structures collected in a folder to be classified based on fold similarity. Based on the results, clusters with structures similar to those of the levan biosynthesis pathway were grouped together.

## 3 Results

The *B. subtilis* strain 168 genome (GenBank Accession ID: GCA_000009045.1), extracted from UniProt using taxonomic identification, had 4191 unique UniProt IDs. But structural data in PDB in relation to UniProt IDs was available for only 753 proteins (Supplementary Table 1). Constraining PDB structures within the range of enzymes present in the GPR association led to 168 unique UniProt IDs with 508 PDB structures. All GPR-associated structures were added to the specific GPR in each reaction. Distinct PDB entries capture complementary mechanistic states. We retained them, prioritizing catalytically representative forms. Some entries were mutants or truncated constructs that might not represent the functional protein. We prioritized catalytically relevant, wild-type, and full-length entries and documented mapping quality for each gene. Each of the 508 PDB structures were assigned to genes labelled in the gene-protein reactions of the *B. subtilis* genome. This provided a direct relationship between structures and reactions. Thus, PDB IDs were assigned for 331 of 1250 reactions (Supplementary Table 2).

**Table 2.** Available PDB structures of the SacB enzyme of *B. subtilis*. These PDB IDs are the structures of levansucrase (SacB) and its mutants.

| PDB ID | Resolution | Chains | Length | Reference |
| --- | --- | --- | --- | --- |
| 1OYG | 1.50 Å | A | 30-473 | (Meng and Fütterer, 2003) |
| 1PT2 | 2.10 Å | A | 30-473 |  |
| 2VDT | 3.20 Å | A | 34-472 | (Ortiz-Soto *et al.*, 2008) |
| 3BYJ | 2.10 Å | A | 1-473 | (Meng and Fütterer, 2008) |
| 3BYK | 2.10 Å | A | 1-473 |  |
| 3BYL | 2.10 Å | A | 1-473 |  |
| 3BYN | 2.10 Å | A | 1-473 |  |
| 6PWQ | 2.60 Å | A/B | 30-473 | (Ortiz-Soto *et al.*, 2020) |
| 6VHQ | 2.05 Å | A/B | 30-473 | (Raga-Carbajal *et al.*, 2021) |

### 3.1 Integration of PDB structures in the levan biosynthesis pathway

We collected structures from the PDB and AlphaFold2 databases. We also identified and oriented membrane-bound proteins by finding transmembrane regions using the TMHMM2.0 server (https://services.healthtech.dtu.dk/services/TMHMM-2.0/). The structural information incorporated into the levan synthesis pathway is represented in Figure 1.

**Figure 1.**
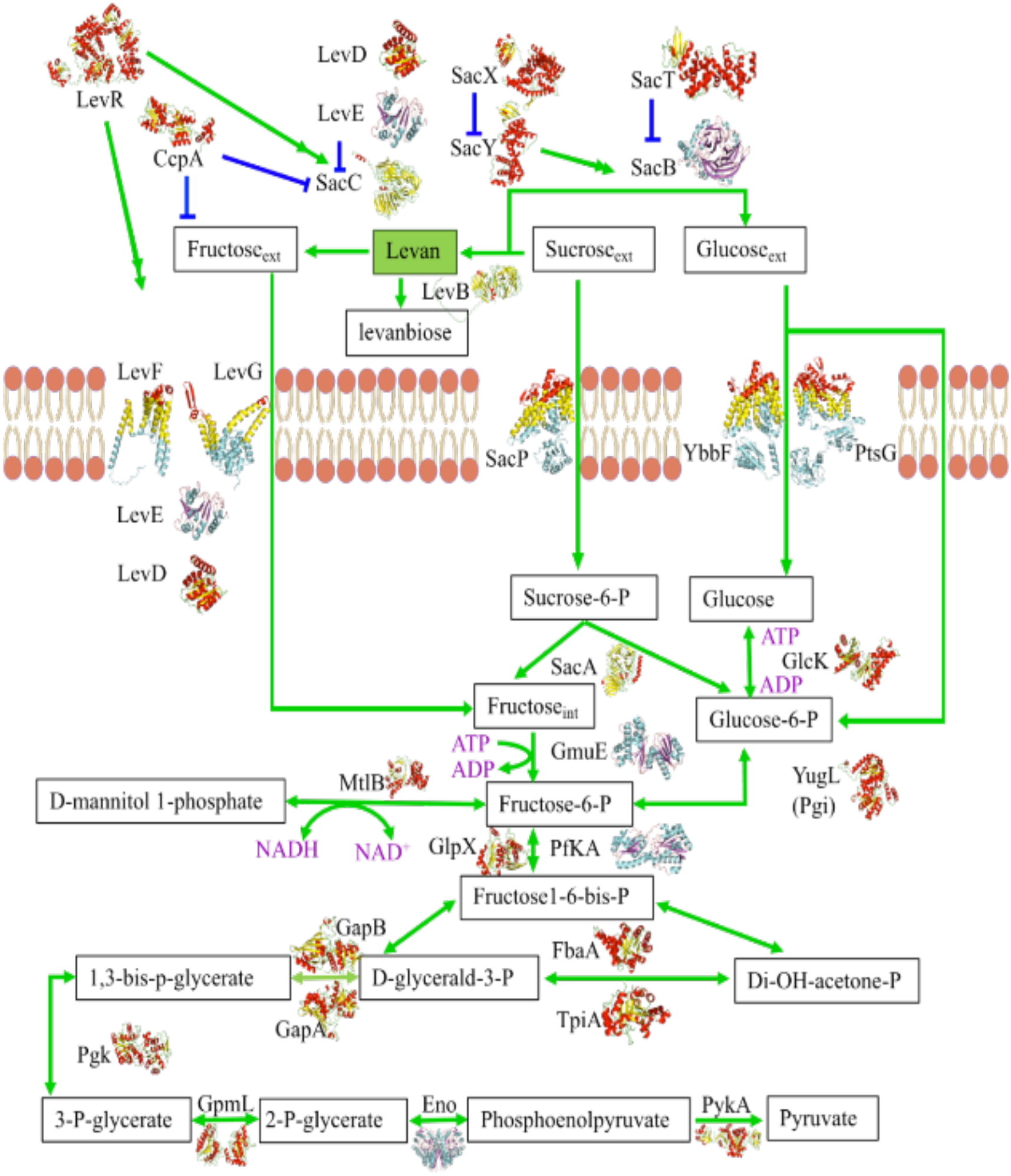
The structural information incorporated into the levan biosynthesis metabolic network. The metabolic network with structural information is provided. Metabolites are represented in a box, and the catalysis of the metabolites is provided in green forward and reversible arrows. Blue arrows indicate regulation information. Structures available in PDB are shown as cyan helices and purple sheets, and AlphaFold2 structures are depicted as yellow sheets and red helices.

Levan biosynthesis is initiated in the absence of glucose as well as in the presence of sucrose. Without glucose, the genes, *sacX* and *sacT*, are deactivated, causing *sacY* to be activated, which, in turn, activates SacB, which produces levan. The levan polymer is then broken down by *levB* into fructan monomer units. These monomer units are transported into the cell by the proteins of the fructose phosphotransferase system (PTS): LevD, LevE, LevF, and LevG. The monomers then join the regular glycolysis pathway to produce energy. The levan pathway is an alternative for the utilization of sucrose. However, sucrose can also be directly transported into the cell by SacP. The transported glucose, sucrose, and fructose contribute towards the growth of the organism.

In the levan biosynthesis pathway, we focused on the SacB enzyme. SacB, involved in the hydrolysis of sucrose and the polymerization of levan in *B. subtilis*, has multiple solved structures (Table 2).

The PDB structures with single point mutations on SacB help understand the mechanism by which the catalytic functional amino acids bind to the substrate and release the product. Mutant studies led to an understanding of the degree of polymerization of levan and to the determination of the polymer’s molecular weight and other aspects. (Meng and Fütterer, 2003, 2008; Ortiz-Soto *et al*., 2008, 2020; Raga-Carbajal *et al*., 2021). Meng and Fütterer identified catalytic residues and amino acids in SacB that are important for its function (1OYG & 1PT2) (Meng and Fütterer., 2008). Ortiz Soto et al. determined amino acids responsible for the degree of polymerization of levan (3BYJ, 3BYK, 3BYL, 3BYN & 6PWQ). Raga Carabajal determined the binding and polymerization mechanism of levan using mutants (6VHQ) (Raga-Carbajal,E. *et al*. 2021). The mutant structures thus provide valuable information for the determination of the functional role of levansucrase.

Other enzymes in the levan biosynthesis/degradation pathway are listed, along with available structures, in Table 3.

**Table 3.** Available PDB structures of levan biosynthesis/degradation and glycolysis pathway enzymes of *B. subtilis*.

| Gene name (aliases) | Gene Identifiers | PDB ID(s) |
| --- | --- | --- |
| *sacB* | BSU34450 | 1OYG;1PT2;2VDT;3BYJ;3BYK;3BYL;3BYN;6PWQ;6VHQ; |
| *sacC* | BSU27030 | 4AZZ;4B1L;4B1M; |
| *sacY (sacS)* | BSU38420 | 1AUU; |
| *ccpA (alsA amyR graR)* | BSU29740 | 1ZVV;2FEP;3OQM;3OQN;3OQO; |
| *ptsG (crr ptsX)* | BSU13890 | 1AX3;1GPR; |
| *levE (sacL)* | BSU27060 | 1BLE; |
| *ptsH* | BSU13900 | 1JEM;1KKL;1KKM;1SPH;2FEP;2HID;2HPR;3OQM;3OQN;3OQO; |
| *gmuE (ydhR)* | BSU05860 | 1XC3;3LM9;3OHR; |
| *pfkA (pfk)* | BSU29190 | 4A3S; |
| *gapB* | BSU29020 | 8WWZ; |
| *eno* | BSU33900 | 4A3R;7XML; |

The permease enzymes IIA and IIB, and the PTS protein enzyme I, a histidine-containing phosphocarrier protein (HPr) are involved in the phosphoenolpyruvate sugar phosphotransferase system, which transfers and phosphorylates a carbohydrate moiety using sugar-specific components. During phosphoryl group transfer, these enzymes form a transient binary complex, initiated by a trigger switch between the salt bridges of the complex of HPr, typically involved in the carbohydrate monomeric unit permease. This mechanistically important alternating ion pair helps in protein-protein phosphotransfer reactions and the transfer of carbohydrates.

PtsG relates to IIA^glc^ and LevE to IIB^lev,^. In PtsG, Arg-17 acts as a switch, while in LevE, His15 acts as a trigger switch. Using the full-length structure of the LevE enzyme, the importance of the residue His15 was identified. LevE is a fructose permease enzyme IIB subunit (IIBLev). LevDEFG transports the fructose moiety, while PtsGHI facilitates the transport of the glucose moiety (Chaptal *et al*., 2006; Chen *et al*., 1998; Fieulaine *et al*., 2002; Herzberg, 1992; Jones *et al*., 1997; Liao and Herzberg, 1994; Schauder *et al*., 1998; Schumacher *et al*., 2011). Notably, most enzymes with the transport mechanism are in the AND class.

### 3.2 Transcriptional regulation using Boolean network

The Boolean network was modeled using BooleSim (Bock *et al*., 2014) (Figure 2) with the gene regulators for levan production in *B. subtilis*. This network visually represents interactions between genes and proteins, showing how they collectively regulate levan production.

**Figure 2.**
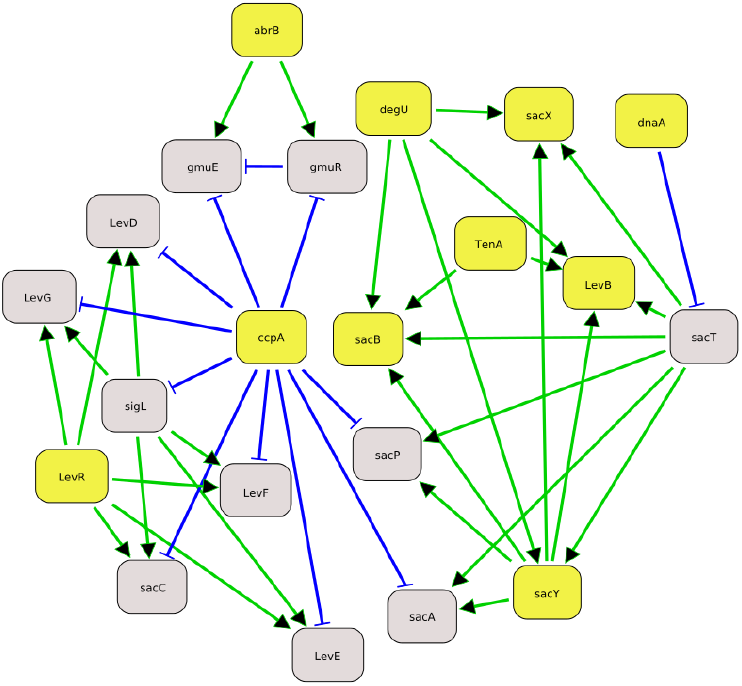
Transcriptional regulatory interaction network of levan biosynthesis. The initial ON and OFF states of genes are shown in lime green and grey, respectively. Green arrows represent activators and blue arrows represent repressors or inhibitors. A blue arrow pointing from one gene to another indicates that the source gene inhibits the target gene. The boxes are nodes. Yellow nodes represent currently active genes involved in cellular processes related to levan production. Gray boxes indicate inactive genes not currently being expressed or contribute to the regulatory network’s activities. The network also allows for a parallel setup where the state of the network can be toggled, switching the colors of the nodes to represent alternate states in the Boolean network. This feature facilitates the visualization of different regulatory scenarios and their impacts on gene expression and levan production.

The Boolean network highlights the dynamic nature of gene regulation in *B. subtilis*. For example, the central node ‘*ccpA*’ plays a crucial role, with multiple outgoing edges indicating its regulatory influence over several genes. Activators like ‘*ccpA*’ can enhance the expression of genes such as ‘*gmuR*’, ‘*levB*’, and ‘*sacX*’, while simultaneously repressing the genes, ‘*sigL*’, ‘*sacB*’, and ‘*levF*’. This intricate balance of activation and repression ensures the fine-tuned regulation of levan production.

By modeling these relationships using BooleSim, we can predict the behavior of the regulatory network under different conditions (True and False states), providing insights into the genetic control mechanisms governing levan production in *B. subtilis*.

The time series simulation in BooleSim was performed to analyze the dynamic behavior of Boolean networks over time. In the simulation, yellow indicates a True state, blue indicates a False state, and the green arrow marks the iterator position. The X-axis represents time in seconds, while the Y-axis represents different genes. This visualization allows us to observe how the states of various genes change over time, providing insights into the temporal dynamics of the regulatory network.

In Figure 3A, the simulation begins with all gene states being True at time t=1. During the simulation, specific genes settle into permanent states: *levD, sigL, levE, sacC, sacT, levF, levG, sacP*, and *sacA* consistently remain False, while *levR, ccpA, degU, levB, sacY, tenA, sacX, sacB*, and *dnaA* consistently remain True. Notably, *abrB, gmuR*, and *gmuE* do not stabilize and oscillate between True and False, indicating the interplay between the three enzymes for which they code and their complex regulatory role in the survival and growth of *B. subtilis*.

**Figure 3.**
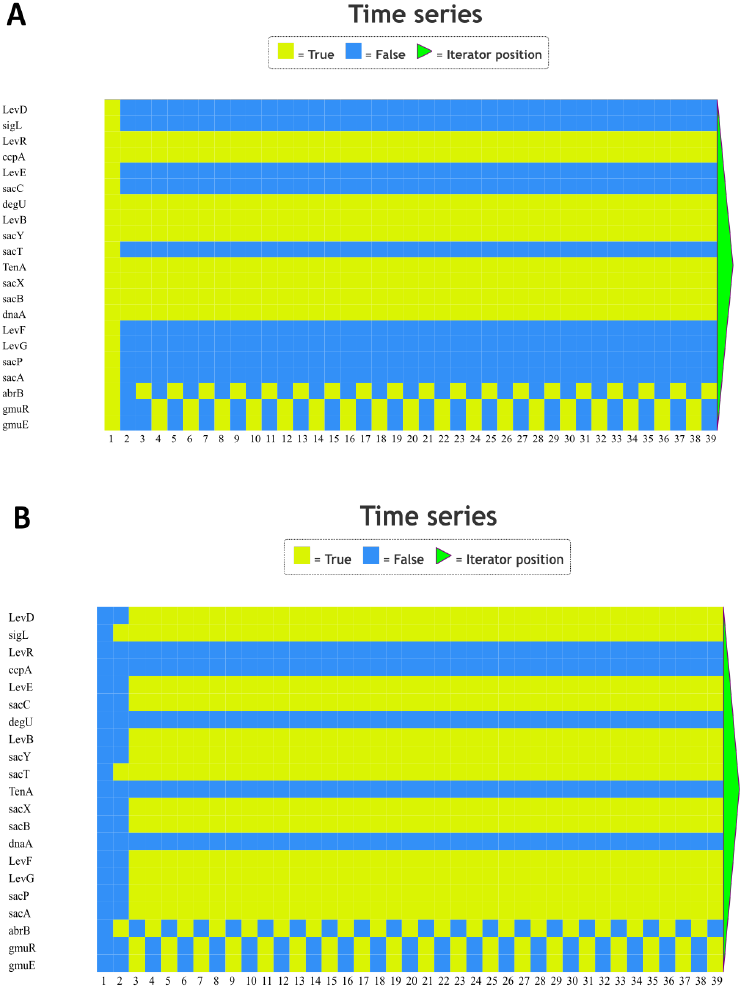
Time series analysis is shown as basic Boolean output functions of 0 and 1. Two initial states were considered: global ON-state 2 (A) and global OFF-state 2 (B). The oscillations allow the network to proceed forward. The ON state is shown as lime green, while the OFF state is shown as light blue.

A different initial condition, with all gene states being False at time t=1 was applied to check the behaviour of the network. This resulted in a distinct pattern of gene states. In the 1st iteration, all the genes were in the OFF or False state. By the 3rd iteration, *levR, ccpA, degU, tenA*, and *dnaA* were in the OFF or False state in all iterations. The remaining genes, *levD, sigL, levE, sacC, sacT, levF, levG, sacP, sacA, levB, sacY, sacX*, and *sacB*, change to the ON or True state in the 3rd iteration. Despite the change in initial conditions, *abrB, gmuR*, and *gmuE* continue to oscillate between the True or ON and False or OFF states, emphasizing their dynamic regulatory roles.

A comparison of Figures 3a and 3b reveals critical insights into the robustness and sensitivity of the gene regulatory network. Specifically, it can be inferred that *levB, sacY, sacX*, and *sacB* are permanently active, irrespective of initial conditions, suggesting that these genes play a crucial and stable role in the network. The consistent behavior of these genes across different simulations underscores their potential importance in maintaining the regulatory balance necessary for levan production.

The fluctuating behavior of *abrB, gmuR*, and *gmuE* under both initial conditions (represented by Figures 3a and 3b) highlights their roles in adaptive responses and conditional regulatory mechanisms within the network. The gene regulatory network of the levan biosynthesis pathway involves two global regulators, ccpA and abrB, within its 1st degree of neighbourhood. Both ccpA and abrB act as repressor, for specific operons involved in carbohydrate utilization (Chumsakul et al, 2011). Since ccpA is turned off under both initial conditions represented in Figure 3, gmuE and gmuR exhibit oscillations in time opposite in phase to that of the repressor, abrB. Thus, by simulating the time series behavior of the Boolean network, BooleSim provided valuable insights into the temporal dynamics and stability of gene regulatory interactions. These findings contribute to a deeper understanding of levan production’s complex regulatory mechanisms in *B. subtilis*, guiding further experimental investigations and potential biotechnological applications.

### 3.3 Integration of structural information into GEM

Newman et al. revealed the relationship between phosphofructokinase (PfkA) and enolase (Eno), where Eno’s product, phosphoenolpyruvate, inhibits the activity of PfkA (Newman *et al*., 2012). The fructokinase enzyme (GmuE) has high specificity towards the fructose moiety and belongs to the ROK glucokinase family. The structures of fructokinase, phosphofructokinase, and enolase were available in the PDB. For the remaining proteins in the pathways for levan biosynthesis and degradation, including glycolysis, AlphaFold2 structures were extracted from the UniProt database and are depicted in the network.

#### 3.3.1 Structure availability-based classification of GPR associations

For further analysis, literature corresponding to PDB IDs was collected and assigned to each GPR. GPR rules, satisfying the conditions for a single protein carrying out the catalysis, were used for further analysis and meaningful interpretation.

#### 3.3.2 Two enzyme OR class

Here, we present some case studies, featuring structural information about the network in the metabolic model. The reaction, catalyzed by the enzymes, BSU09530 and BSU39780, corresponds to two aldo-keto reductases (AKRs), AKR11A and AKR11B, in *B. subtilis*. AKR11B has the conventional structure of AKRs and is more efficient than AKR11A. AKR11A is speculated to have a different primary role from that of an oxidoreductase. Due to the masking effects of AKRs, it is safe to knock out AKR11A to allow the efficient AKR11B enzyme to benefit from oxidoreductase activity (Ehrensberger and Wilson, 2004).

While YvgN belongs to AKR5G1 and YtbE to AKR5G2, there is another set of AKRs in *B. subtilis* involved in detoxifying various aldehydes generated by stress. These AKRs catalyze the reduction of an aldose or aldehyde substrate to an alcohol using NADPH. These proteins, with 70% similarity, have distinct differences in the C-terminal α/β barrel, especially in the loops, β1 and β7. The loops are connected by hydrogen bonds, forming a safety belt stabilized by the K26 residue in YvgN and by the G26 residue in YtbE. Both proteins are essential for survival and growth. While YtbE is an intracellular protein, YvgN is extracellular. (Lei *et al*., 2009). This could perhaps explain the differences in structural topology.

Similarly, in the case of dUTPases, *B. subtilis* contains a prophage dUTPase and the genomic dUTPase. These two homotrimeric enzymes catalyze the hydrolysis of 2-deoxy-uridine triphosphate (dUTP) to 2-deoxy-uridine monophosphate (dUMP) in an ion-dependent process. Both enzymes are structurally different from regular dUTPases. They have a ‘Phe-lid’ on the uracil moiety, whereas, in regular dUTPases, motif V is conserved. Consequently, the catalytic mechanism is different. These two proteins are similar at the sequence level, which suggests horizontal gene transfer. Similar evolutionary patterns have been detected in dUTPase enzymes in other organisms (García-Nafría *et al*., 2010, 2011, 2013).

In Gram-positive bacteria, such as *B. subtilis*, class D β-lactamases have an extended loop1 region in β8 and β9, forming a hydrophobic cap over the active site. Deleting this loop region significantly increases the catalytic activity of the enzyme.

Glutamate metabolism in *B. subtilis* is catalyzed by the glutamate dehydrogenases, RocG and GudB. The GudB enzyme is cryptic due to the insertion of three residues near the active site. In the absence of RocG, GudB replaces the function. It also inhibits the synthesis of glutamate by binding to glutamate synthase (GltAB), playing a dual role in glutamate homeostasis (Gunka *et al*., 2010; Jayaraman *et al*., 2022).

The ureide pathway, essential for transporting and storing nitrogen, involves the hydrolysis of uric acid into ureides using the enzymes urate oxidase (BsUOX) and PucM. Though structurally unrelated, these two proteins play a role in the same metabolic pathway due to their similar functions (Nayab *et al*., 2019). The PucM protein, 5-hydroxyisourate hydrolase, essential for the purine degradation pathway, is highly sensitive to mutation and small changes in the sequence lead to functional differences (Jung *et al*., 2006).

#### 3.3.3 Two enzyme AND class

The enzymes of the two enzyme AND class catalyze a particular reaction step, using different approaches, such as complex formation, signaling and interaction. Unlike the OR function, the AND function is very sparse. Structures were only available for YaaE and YaaD, the enzymes involved in vitamin B6 biosynthesis. The enzymes, YaaE and YaaD, form an oligomeric barrel structure. The complex is directed by the N-terminal α-helix, which precedes glutaminase activation, forming an oxyanion hole. This oxyanion hole facilitates the interaction of the ammonia molecule with the YaaD active site (Bauer *et al*., 2004; Strohmeier *et al*., 2006).

#### 3.3.4 Multi-enzyme OR group

From the structural biology perspective, chorismate mutase is a well-studied protein in *B. subtilis*. The protein is involved in the aromatic amino acid biosynthetic pathway and catalyzes the conversion of chorismate to prephenate with a single transition state between the reactions. Three different enzymes possess the ability to perform a similar function with more detailed structural data present for AroQ (BSU22690) and AroH (BSU29750) (Burschowsky *et al*., 2014; Chook *et al*., 1994, 1994; Kast *et al*., 2000; Ladner *et al*., 2000; Pratap *et al*., 2017). These studies suggest that the enzyme, AroQ, has lower catalytic efficiency. AroQ facilitates the capture of important enzyme catalysis origins, as explained by the transition state theory, a paradigm proposed by Pauling (Pauling, 1946). The transition state theory states that enzymes bind to transition states more tightly than substrates, thereby lowering activation energy and accelerating the chemical reaction. Structural studies for AroQ showed that the shape of the active site, complementary to the substrate, is not a major factor for efficient catalysis. However, the electrostatic organization of the native substrate and electrostatic stabilization during the transition state at the active site are important to achieve high reaction rates.

### 3.4 Analysis of structure-based clustering

We proceeded with available AlphaFold2 structures from the Uniprot and AlphaFold databases. There were eight protein sequences for which no full-length structures were available in either the Uniprot or the AlphaFold database.

These were sequences with the Uniprot IDs: O07005, O31782, P27206, P40806, P40872, P94459, Q04747, and Q05470. Thus, for clustering analysis, 4183 structures were used.

Clustering analysis based on AlphaFold2 structures using FoldSeek showed clusters of protein folds that were conserved; these structures were classified based on fold architecture. There were 563 unique folds, with 1589 structures not clustered among known fold architectures. Levan biosynthesis enzymes were clustered into four groups as shown in Figure 4, sharing the same fold architecture as other proteins in *B. subtilis*.

**Figure 4.**
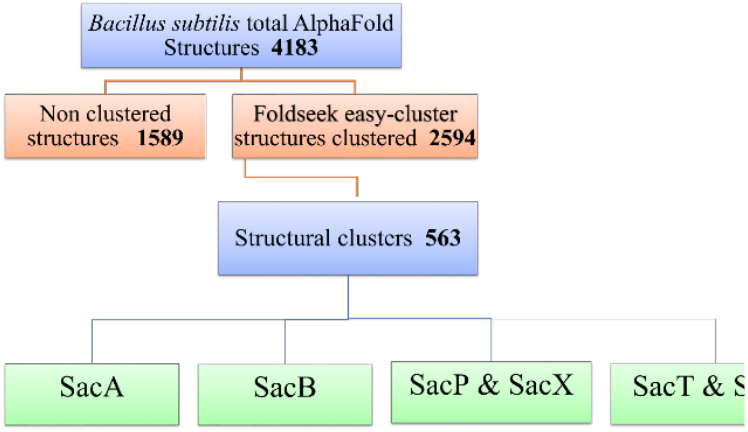
Clustering analysis of 4183 AlphaFold2 structures of the *B. subtilis* proteome. Matching folds were found in 2594 structures. They were clustered into 563 cluster-based matches depending on fold level information. Unique folds that were not part of any cluster were found in 1589 structures. Levan-associated enzymes were separated into four different clusters.

As given in Supplementary Table 1 and Figure 5, the structural architectures of SacB and SacA were clustered and classified as carbohydrate-associated enzymes. SacB, consisting of 10 proteins, was classified as an amino acid biosynthetic cluster using carbohydrates as precursors. The structure of GH68, one of the proteins in the SacB cluster, consistently showed a five-bladed β-propeller conformation. The folds feature five β-sheet blades organized around a central axis, creating a single-domain structure that encases a funnel-shaped central cavity containing the active site (Veerapandian *et al*., 2025).

**Figure 5.**
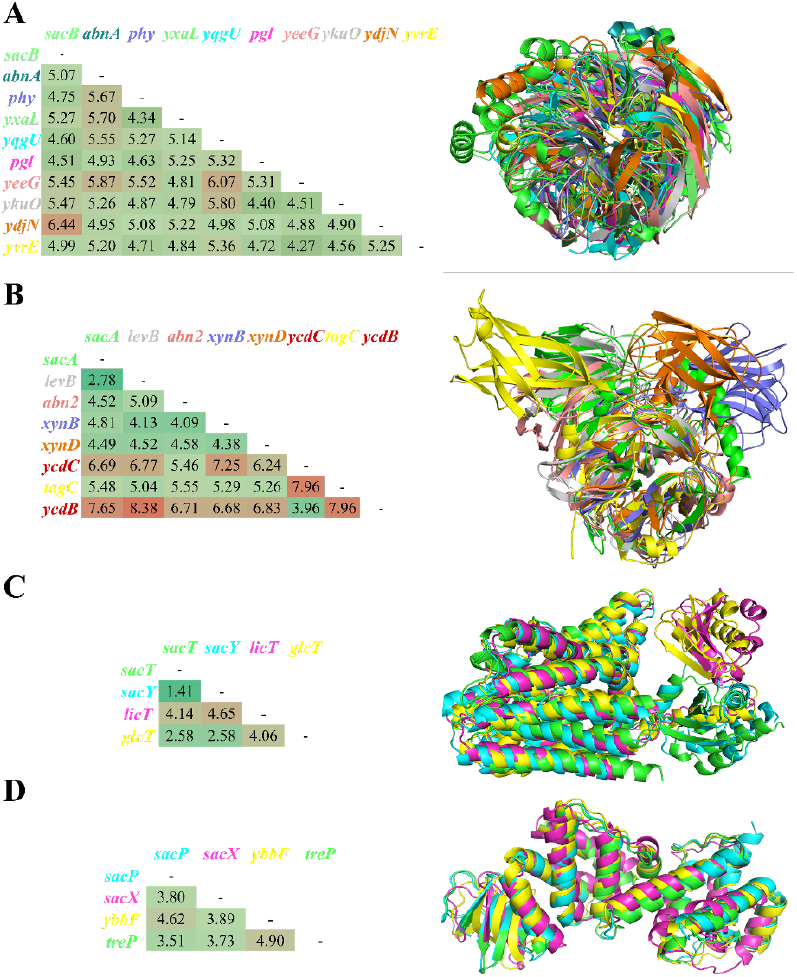
Structural superposition-based comparison of levan-associated structural clusters. The representative structure (green) superposed with cluster structures in other colors. RMSD correlation is shown on the sides of the superposed structures. A) The SacB cluster contains the glycoside hydrolase family 68 (GH68) fold, and the results revealed that several other proteins in B. subtilis share a similar fold. B) The SacA cluster shows higher RMSD values, and the fold architecture of glycoside hydrolase family 32 (GH32) is conserved. C) SacP and SacX share a similar fold within the levan biosynthesis enzyme. D) SacT and SacY share similar architecture with low RMSD deviation amongst all levan biosynthesis clusters. The RMSD cross-correlation matrix is represented adjacent to each superimposed cluster; each structure is represented based on its specific color.

SacA, consisting of eight proteins, was classified as a cluster involved in the carbohydrate catabolic process and cell wall biosynthesis. SacT and SacY were clustered together with four other proteins and were classified as positive regulators of DNA transcription. Similarly, SacP and SacX were clustered together with four other proteins and were classified as a phosphoenolpyruvate-dependent sugar phosphotransferase system.

We were able to identify and classify seven uncharacterized proteins within levan biosynthesis proteins with similar fold and domain architecture. The functions of these seven uncharacterized proteins need to be explored via experimental methods to appropriately annotate the *B. subtilis* genome.

## 4 Conclusion

In this study, we brought experimentally derived and computationally predicted structures together into the levan biosynthesis pathway of *B. subtilis*. Integrating the PDB structures into the GEM provided detailed structural insights into the metabolic network. With the help of a Boolean gene regulatory network and time series analysis, we delineated the interdependence of the genes within the levan biosynthesis pathway. The integration of the gene regulatory network into the levan biosynthesis pathway model will significantly enhance prediction accuracy, improving our ability to predict levan production in *B. subtilis*.

We quantified structural coverage across the model: 331 out of 1250 reactions (26.5%) and 508 PDB entries. We also identified 563 clusters based on the folding characteristics among the predicted AlphaFold2 structures of the *B. subtilis* proteome. Similarities in folding helped group proteins involved in the levan biosynthesis pathway into four distinct clusters. Though the proteins in each cluster did not show sequence similarity, our results suggest that each cluster has distinct functions. Clustering also facilitated the classification of seven uncharacterized proteins of unknown function in the levan biosynthesis pathway.

The comprehensive approach outlined in this study will advance the understanding of the molecular and regulatory mechanisms underlying levan biosynthesis and provide a robust framework for optimizing metabolic processes with structural data in *B. subtilis*, expanding the capabilities of genome-scale metabolic models.

## Acknowledgements

Dr. UV acknowledges DST SERB-CRG (CRG/2023/006281). The authors thank SASTRA Deemed to be University for the infrastructural support and financial support through the Research and Modernization funding.

## Supplementary data

Supplementary data are available at *Bioinformatics Advances* online.

## Data availability

The data underlying this article are available in: *levan_sssbio* at https://github.com/raghuyennamalli/levan_ssbio

