## Supplementary Tables for "Structural and Boolean Network Modeling of the Levan Biosynthetic Pathway in *Bacillus subtilis*"

**Supplementary Material**

**List of Supplementary Tables**

| **S.No.** | **Contents** | **Page No.** |
| --- | --- | --- |
| **1** | **List of clusters of structurally similar folded proteins according to FoldSeek.** | **1** |
| **2** | **Available PDB structures for *B. subtilis*** | **4** |
| **3** | **GPR-PDB structure associations** | **38** |

**Supplementary Table 1.** List of clusters of structurally similar folded proteins according to FoldSeek.

| **Cluster** | **UniProt ID** | **Gene ID** | **Gene Name** | **GO: Biological Process** | **CAZy family** | **E.C. Number** | **Enzyme Name** | **Subject /Query**  **Seqeunce Identity** |
| --- | --- | --- | --- | --- | --- | --- | --- | --- |
| SacB | P05655 | BSU34450 | *sacB* | Carbohydrate utilization | GH68 - Glycoside Hydrolase Family 68 | 2.4.1.10 | Levansucrase | 473/473 (100.0%) |
|  | P94522 | BSU28810 | *abnA* | Arabinan catabolic process | GH43 - Glycoside Hydrolase Family 43 | 3.2.1.99 | Extracellular endo-alpha-(1->5)-L-arabinanase 1 | 94/499 (18.8%) |
|  | P42094 | BSU19800 | *phy* | - | - | 3.1.3.8 | 3-phytase | 97/538 (18.0%) |
|  | P42111 | BSU39940 | *yxaL* | - | - | - | Uncharacterized protein | 84/577 (14.6%) |
|  | P54498 | BSU24820 | *yqgU* | - | - | - | Uncharacterized lipoprotein | 78/538 (14.5%) |
|  | O34499 | BSU13010 | *pgl* | Glucose metabolic process, pentose-phosphate shunt | - | 3.1.1.31 | 6-phosphogluconolactonase | 79/559 (14.1%) |
|  | O31507 | BSU06820 | *yeeG* | - | - | - | Uncharacterized protein | 80/573 (14.0%) |
|  | O34879 | BSU14160 | *ykuO* | - | - | - | Uncharacterized protein | 70/516 (13.6%) |
|  | O34353 | BSU06260 | *ydjN* | - | - | - | Uncharacterized protein | 73/561 (13.0%) |
|  | O34940 | BSU33200 | *yvrE* | L-ascorbic acid biosynthetic process | - | 3.1.1.- | Putative sugar lactone lactonase | 54/563 ( 9.6%) |
| SacA | P07819 | BSU38040 | *sacA* | Sucrose metabolic process | GH32 - Glycoside Hydrolase Family 32 | 3.2.1.26 | Sucrose-6-phosphate hydrolase | 479/479 (100.0%) |
|  | O07003 | BSU34460 | *levB* | Sucrose catabolic process | GH32 - Glycoside Hydrolase Family 32 | 3.2.1.64 | Levanbiose-producing levanase | 134/589 (22.8%) |
|  | P42293 | BSU39330 | *abn2* | Arabinan catabolic process | GH43 - Glycoside Hydrolase Family 43 | 3.2.1.99 | Extracellular endo-alpha-(1->5)-L-arabinanase 2 | 95/592 (16.0%) |
|  | P94489 | BSU17580 | *xynB* | Xylan catabolic process | GH43 - Glycoside Hydrolase Family 43 | 3.2.1.37 | Beta-xylosidase | 101/645 (15.7%) |
|  | Q45071 | BSU18160 | *xynD* | Xylan catabolic process | CBM6 - Carbohydrate-Binding Module Family 6  GH43 - Glycoside Hydrolase Family 43 | 3.2.1.55 | Arabinoxylan arabinofuranohydrolase | 95/666 (14.3%) |
|  | O34772 | BSU02800 | *ycdC* | - | - | - | Uncharacterized protein | 67/722 ( 9.3%) |
|  | P27622 | BSU35770 | *tagC* | Cell wall organization, teichoic acid biosynthetic process | - | - | Putative major teichoic acid biosynthesis protein C | 62/715 ( 8.7%) |
|  | O34621 | BSU02790 | *ycdB* | - | - | - | Uncharacterized protein | 26/867 ( 3.0%) |
| SacP & SacX | P05306 | BSU38050 | *sacP* | Phosphoenolpyruvate-dependent sugar phosphotransferase system,  trehalose transport | - | 2.7.1.211 | PTS system sucrose-specific EIIBC component | 461/461 (100.0%) |
|  | P15400 | BSU38410 | *sacX* | Phosphoenolpyruvate-dependent sugar phosphotransferase system,  trehalose transport | - | 2.7.1.- | Probable PTS system sucrose-specific EIIBC component | 260/463 (56.2%) |
|  | P39794 | BSU07800 | *treP* | Phosphoenolpyruvate-dependent sugar phosphotransferase system,  trehalose transport | - | 2.7.1.201 | PTS system trehalose-specific EIIBC component | 194/486 (39.9%) |
|  | Q797S1 | BSU01680 | *ybbF* | Phosphoenolpyruvate-dependent sugar phosphotransferase system | - | 2.7.1.- | Putative PTS system EIIBC component | 171/478 (35.8%) |
| SacT & SacY | P26212 | BSU38070 | *sacT* | Positive regulation of DNA-templated transcription | - | - | SacPA operon antiterminator | 276/276 (100.0%) |
|  | P15401 | BSU38420 | *sacY* | Positive regulation of DNA-templated transcription | - | - | Levansucrase and sucrase synthesis operon antiterminator | 133/280 (47.5%) |
|  | P39805 | BSU39080 | *licT* | Positive regulation of DNA-templated transcription | - | - | Transcription antiterminator | 113/279 (40.5%) |
|  | O31691 | BSU13880 | *glcT* | Positive regulation of DNA-templated transcription | - | - | PtsGHI operon antiterminator | 94/283 (33.2%) |

**Supplementary Table 2.** Available PDB structures for *B. subtilis.* The gene name, as well as alias and gene identifier corresponding to its PDB id are provided

| **Gene Name & Identifier** | **PDB ID** |
| --- | --- |
| pyrR BSU15470 | 1A3C;1A4X;4P82; |
| rnpA BSU41050 | 1A6F;4JG4; |
| cpfC hemF hemH BSU10130 | 1AK1;1C1H;1C9E;1DOZ;1LD3;1N0I;2AC2;2AC4;2H1V;2H1W;2HK6;2Q2N;2Q2O;2Q3J;3GOQ;3M4Z; |
| purF BSU06490 | 1AO0;1GPH; |
| sacY sacS BSU38420 ipa-13r | 1AUU; |
| spoIIAA BSU23470 | 1AUZ;1BUZ; |
| ptsG crr ptsX BSU13890 | 1AX3;1GPR; |
| xynA BSU18840 | 1AXK;1XXN;2B42;2B45;2B46;2DCY;2DCZ;2QZ3;2Z79;3EXU;3HD8;5K9Y;5TVV;5TVY;5TZO; |
| thyA1 BSU17680 | 1B02;1BKO;1BKP;1BSF;1BSP; |
| sinR flaD sin BSU24610 | 1B0N;2YAL;3QQ6;3ZKC;5TN0;5TN2; |
| sinI BSU24600 | 1B0N;5TMX; |
| amyE amyA BSU03040 | 1BAG;1UA7; |
| levE sacL BSU27060 | 1BLE; |
| rtp BSU18490 | 1BM9;1F4K;1J0R;2DPD;2DPU;2DQR;2EFW; |
| pel BSU07560 | 1BN8;2BSP;2NZM;2O04;2O0V;2O0W;2O17;2O1D;3KRG;5AMV;5X2I; |
| bmrR bmr1R BSU24020 | 1BOW;1EXI;1EXJ;1R8E;2BOW;3D6Y;3D6Z;3D70;3D71;3IAO;3Q1M;3Q2Y;3Q3D;3Q5P;3Q5R;3Q5S;7CKQ; |
| thiM thiK ywbJ BSU38300 ipa-25d | 1C3Q;1EKK;1EKQ;1ESJ;1ESQ; |
| pnbA estB BSU34390 | 1C7I;1C7J;1QE3; |
| aroH BSU22690 | 1COM;1DBF;1FNJ;1FNK;2CHS;2CHT;3ZO8;3ZOP;3ZP4;3ZP7; |
| tagD BSU35740 | 1COZ;1N1D; |
| cspB cspA BSU09100 | 1CSP;1CSQ;1NMF;1NMG;2ES2;2F52;2I5L;2I5M;3PF4;3PF5;6SZZ;6T00; |
| pyrF BSU15550 | 1DBT; |
| prs BSU00510 | 1DKR;1DKU;1IBS; |
| nadE outB BSU03130 | 1EE1;1FYD;1IFX;1IH8;1KQP;1NSY;2NSY; |
| maf BSU28050 | 1EX2;1EXC;4HEB; |
| purB purE BSU06440 | 1F1O; |
| spo0F BSU37130 | 1F51;1FSP;1NAT;1PEY;1PUX;1SRR;2FSP;2FTK;2JVI;2JVJ;2JVK;3Q15; |
| spo0B spo0D BSU27930 | 1F51;1IXM;2FTK; |
| acpS ydcB BSU04620 | 1F7L;1F7T;1F80; |
| acpA acpP BSU15920 | 1F80;1HY8;2X2B; |
| argR ahrC BSU24250 | 1F9N;2P5K;2P5L;2P5M; |
| gerE BSU28410 | 1FSE; |
| thiE thiC ywbK BSU38290 ipa-26d | 1G4E;1G4P;1G4S;1G4T;1G67;1G69;1G6C;2TPS;3O15;3O16; |
| yqhS BSU24470 | 1GQO; |
| cotA pig BSU06300 | 1GSK;1OF0;1W6L;1W6W;1W8E;2BHF;2WSD;2X87;2X88;3ZDW;4A66;4A67;4A68;4AKO;4AKP;4AKQ;4Q89;4Q8B;4YVN;4YVU;5ZLJ;5ZLK;5ZLL;5ZLM;7Y8B;7Y8C; |
| spsA BSU37910 ipa-63d | 1H7L;1H7Q;1QG8;1QGQ;1QGS; |
| licT BSU39080 N15A | 1H99;1L1C;1TLV;6TWR; |
| dppA dciAA BSU12920 | 1HI9; |
| icd citC BSU29130 | 1HQS; |
| iolI yxdH BSU39680 B65B | 1I60;1I6N; |
| estA lip lipA BSU02700 | 1I6W;1ISP;1R4Z;1R50;1T2N;1T4M;2QXT;2QXU;3D2A;3D2B;3D2C;3QMM;3QZU;5CRI;5CT4;5CT5;5CT6;5CT8;5CT9;5CTA;5CUR; |
| luxS ytjB BSU30670 | 1IE0;1J98;1JQW;1JVI;1YCL;2FQO;2FQT; |
| speE ywhF BSU37500 | 1IY9; |
| cypC CYP152A1 BSU02100 | 1IZO;2ZQJ;2ZQX;7WYG; |
| oxdC yvrK BSU33240 | 1J58;1L3J;1UW8;2UY8;2UY9;2UYA;2UYB;2V09;3S0M;4MET;5HI0;5VG3;6TZP;6UFI; |
| mta ywnD BSU36600 | 1JBG;1R8D; |
| ptsH BSU13900 | 1JEM;1KKL;1KKM;1SPH;2FEP;2HID;2HPR;3OQM;3OQN;3OQO; |
| arsC yqcM BSU25780 | 1JL3;1Z2D;1Z2E;2IPA; |
| srfAC srfA3 BSU03510 | 1JMK;2VSQ;8F7F;8F7G;8F7H;8F7I; |
| ykfB BSU12980 | 1JPM;1TKK; |
| cdd BSU25300 | 1JTK;1UWZ;1UX0;1UX1; |
| copA yvgX BSU33500 | 1JWW;1KQK;1OPZ;1OQ3;1OQ6;1P6T;2RML;2VOY; |
| copZ yvgY BSU33510 | 1K0V;1P8G;2QIF;3I9Z; |
| crh yvcM BSU34740 | 1K1C;1MO1;1MU4;1ZVV;2AK7;2RLZ; |
| ppaC BSU40550 | 1K23;1WPM;1WPN;2HAW;2IW4; |
| nadD yqeJ BSU25640 | 1KAM;1KAQ; |
| nnrD yxkO BSU38720 | 1KYH;3RPH;3RPZ;3RQ2;3RQ5;3RQ6;3RQ8;3RQH;3RQQ;3RQX; |
| cah BSU03180 | 1L7A;1ODS;1ODT; |
| obg BSU27920 | 1LNZ; |
| spo0A spo0C spo0G BSU24220 | 1LQ1; |
| ktrA yuaA BSU31090 | 1LSU;2HMS;2HMT;2HMU;2HMV;2HMW;4J7C;4J90;4J91;5BUT;6S2J;6S5B;6S5C;6S5D;6S5E;6S5G;6S5N;6S5O;6S7R;8K16;8K1K;8K1S;8K1T;8K1U;8XMH;8XMI; |
| hxlB yckF BSU03450 | 1M3S;1VIV; |
| secA div+ BSU35300 | 1M6N;1M74;1TF2;1TF5;2IBM;3DL8;3IQM;3IQY;3JV2;5EUL;6ITC;7XHA;7XHB; |
| nos yflM BSU07630 | 1M7V;1M7Z;2AMO;2AN0;2AN2;2FBZ;2FC1;2FC2;4D3I;4D3J;4D3K;4D3M;4D3N;4D3O;4D3T;4D3U;4D3V;4D7H;4D7I;4D7J;4LWA;4LWB;4UG5;4UG6;4UG7;4UG8;4UG9;4UGA;4UGB;4UGC;4UGD;4UGE;4UGF;4UGG;4UGH;4UGI;4UGJ;4UGK;4UGL;4UGM;4UGN;4UGO;4UGP;4UGQ;4UGR;4UGS;4UGT;4UGU;4UGV;4UGW;4UGX;4UGY;4UQR;4UQS;5G65;5G66;5G67;5G68;5G69;5G6A;5G6B;5G6C;5G6D;5G6E;5G6F;5G6G;5G6H;5G6I;5G6J;5G6K;5G6L;5G6M;5G6N;5G6O;5G6P;5G6Q;6XCX;6XK3;6XK4;6XK5;6XK6;6XK7;6XK8;6XMC; |
| dhbE entE BSU31980 | 1MD9;1MDB;1MDF; |
| yqjY BSU23690 | 1MK4; |
| glsA1 glsA ybgJ BSU02430 | 1MKI;2OSU;3AGF;3BRM; |
| phoP BSU29110 | 1MVO; |
| yteR BSU30120 | 1NC5;2D8L;2GH4; |
| ndoA mazF ydcE BSU04660 | 1NE8;4MDX;4ME7; |
| thiO goxB yjbR BSU11670 | 1NG3;1NG4;1RYI;3IF9; |
| yqeY BSU25400 | 1NG6; |
| yojF BSU19470 | 1NJH; |
| azr yhdA BSU09340 | 1NNI;2GSW;3GFQ;3GFR;3GFS; |
| ywpJ BSU36290 | 1NRW; |
| ydaF BSU04210 | 1NSL; |
| pdaA yfjS BSU07980 | 1NY1;1W17;1W1A;1W1B; |
| purR yabI BSU00470 | 1O57;1P4A;7RMW; |
| coaD ylbI BSU15020 | 1O6B; |
| mnaA BSU35660 | 1O6C;4FKZ; |
| rbsD BSU35930 | 1OGC;1OGD;1OGE;1OGF; |
| yycN BSU40290 | 1ON0;1UFH; |
| mntR yqhN BSU24520 | 1ON1;1ON2;2EV0;2EV5;2EV6;2F5C;2F5D;2F5E;2F5F;2HYF;2HYG;3R60;3R61;4HV5;4HV6;4HX4;4HX7;4HX8;9C4C;9C4D; |
| ypmQ BSU21750 | 1ON4;1XZO; |
| yesU BSU07030 | 1OQ1; |
| hemAT yhfV BSU10380 | 1OR4;1OR6; |
| yuaD BSU31040 | 1ORU; |
| sacB BSU34450 | 1OYG;1PT2;2VDT;3BYJ;3BYK;3BYL;3BYN;6PWQ;6VHQ; |
| rph BSU28370 | 1OYP;1OYR;1OYS; |
| fni idi ypgA BSU22870 | 1P0K;1P0N; |
| adk BSU01370 | 1P3J;2EU8;2OO7;2ORI;2OSB;2P3S;2QAJ;3DKV;3DL0;4MKF;4MKG;4MKH;4QBF;4QBG;4TYP;4TYQ;5X6I; |
| rbgA ylqF BSU16050 | 1PUJ;6PPK; |
| sda BSU25690 | 1PV0;3FYR; |
| ywqG BSU36220 | 1PV5; |
| sboA sbo BSU37350 | 1PXQ; |
| iolS yxbF BSU39780 SS92ER | 1PYF;1PZ0; |
| yhdN BSU09530 | 1PZ1; |
| yjcF BSU11840 | 1Q2Y; |
| yjcS BSU11970 | 1Q8B; |
| ridA yabJ BSU00480 | 1QD9; |
| sfp lpa-8 BSU03570 | 1QR0;4MRT; |
| yvyI pmi BSU35790 | 1QWR; |
| ywiB BSU37340 | 1R0U; |
| pdxT yaaE BSU00120 | 1R9G;2NV0;2NV2; |
| yfiR BSU08370 | 1RKT; |
| ywqN BSU36150 | 1RLI; |
| nrdI ymaA BSU17370 | 1RLJ; |
| yvqK BSU33150 | 1RTY; |
| ribH BSU23250 | 1RVV;1ZIS; |
| yfiT BSU08390 | 1RXQ; |
| recU prfA yppB BSU22310 | 1RZN;1ZP7;5FDK; |
| mdtR yusO BSU32870 | 1S3J; |
| yojM BSU19400 | 1S4I;1U3N;1XTL;1XTM; |
| yesE yeeL BSU06870 | 1S5A; |
| ykoF BSU13240 | 1S7H;1S99;1SBR; |
| aprE apr aprA sprE BSU10300 | 1SCJ;3WHI;6O44;6PAK; |
| yhaI BSU09980 | 1SED; |
| yfhH BSU08530 | 1SF9; |
| yxaF BSU39990 S14F | 1SGM; |
| resA ypxA BSU23150 | 1ST9;1SU9;2F9S;2H19;2H1A;2H1B;2H1D;2H1G;3C71;3C73; |
| engB ysxC BSU28190 | 1SUL;1SVI;1SVW; |
| yvdD BSU34640 | 1T35; |
| csrA sow yviG BSU35370 | 1T3O; |
| purS yexA BSU06460 | 1T4A;1TWJ; |
| iolA mmsA yxdA BSU39760 E83A | 1T90; |
| rsgA cpgA engC yloQ BSU15780 | 1T9H; |
| pta ywfJ BSU37660 ipa-88d | 1TD9;1XCO; |
| paiA BSU32150 | 1TIQ; |
| guaD gde BSU13170 | 1TIY;1WKQ; |
| ypjQ BSU21830 | 1TLQ; |
| rlmH yydA BSU40230 | 1TO0; |
| tenA BSU11650 | 1TO9;1TYH;1YAF;1YAK;2QCX; |
| yhfP BSU10320 | 1TT7;1Y9E; |
| cmoJ moxC ytnJ BSU29310 | 1TVL;1YW1;6ASK;6ASL; |
| yycE BSU40430 | 1TWU; |
| thiS yjbS BSU11680 | 1TYG; |
| thiG yjbT BSU11690 | 1TYG;1XM3; |
| yitD BSU10950 | 1U83; |
| glvA glv-1 glvG malA BSU08180 | 1U8X; |
| abnA BSU28810 | 1UV4; |
| yjbI BSU11560 | 1UX8; |
| ydeN BSU05260 | 1UXO; |
| hutP BSU39340 | 1VEA;1WMQ;1WPS;1WPT;1WPU;1WPV;1WRN;1WRO;1WRQ;3BOY;4H4L; |
| fliS BSU35330 | 1VH6;5MAW;6GOW; |
| ysdC BSU28820 | 1VHE; |
| rsmE yqeU BSU25440 | 1VHK; |
| ywnH BSU36560 | 1VHS; |
| yrrK yqgf BSU27390 | 1VHX; |
| fadR ysiA BSU28550 | 1VI0;3WHB;3WHC; |
| plsX ylpD BSU15890 | 1VI1;6A1K; |
| pcrB yerE BSU06600 | 1VIZ;3VZX;3VZY;3VZZ;3W00; |
| spsE BSU37870 ipa-67d | 1VLI; |
| yxbC yxaQ BSU39880 VE7D | 1VRB;7ZCC; |
| hslO yacC BSU00710 | 1VZY; |
| rsbU BSU04700 | 1W53;2J6Y;2J6Z;2J70; |
| dacC pbp BSU18350 | 1W5D;2J9P; |
| mtrB BSU22770 | 1WAP;3ZZQ;4B27; |
| rsbQ yvfQ BSU34100 | 1WOM;1WPR; |
| lon1 lonA BSU28200 | 1X37;3M65;3M6A; |
| cwlC BSU17410 | 1X60; |
| gmuE ydhR BSU05860 | 1XC3;3LM9;3OHR; |
| ywnA BSU36630 | 1XD7; |
| rsmG gidB BSU41000 | 1XDZ; |
| sufU iscU nifU yurV BSU32680 | 1XJS;2AZH;5XT5;5XT6;6JZV;6JZW; |
| yqbG BSU26120 | 1XN8;1ZTS; |
| appA BSU11381/BSU11382 BSU11380 | 1XOC; |
| ycgJ BSU03160 | 1XXL;2GLU; |
| xpt BSU22070 | 1Y0B;2FXV;6W1I; |
| hit yhaE BSU10030 | 1Y23; |
| qdoI yxaG BSU39980 S14G | 1Y3T;2H0V;8HFB; |
| rnz yqjK BSU23840 | 1Y44;2FK6;4GCW; |
| ykuD BSU14040 | 1Y7M;2MTZ;3ZQD;4A1I;4A1J;4A1K;4A52; |
| tenI BSU11660 | 1YAD;3QH2; |
| darB ykuL BSU14130 | 1YAV;6YJ7;6YJ8;6YJ9;6YJA;8ACU;8AD6; |
| yqgN BSU24890 | 1YDM; |
| yngG BSU18230 | 1YDO; |
| abrB cpsX BSU00370 | 1YFB;1YSF;1Z0R;2K1N;2MJG;2RO4; |
| mstX BSU31321 BSU31320 | 1YGM; |
| xynB BSU17580 | 1YIF; |
| yutE hepT BSU32300 | 1YLM; |
| scmP sndB yxeP BSU39470 LP9H | 1YSJ; |
| yvbK BSU33890 | 1YVK; |
| ysnE BSU28330 | 1YX0; |
| gcvT yqhI BSU24570 | 1YX2; |
| queA BSU27720 | 1YY3; |
| clpQ codW hslV BSU16150 | 1YYF;2Z3A;2Z3B; |
| spx spxA yjbD BSU11500 | 1Z3E;3GFK;3IHQ;6GHB;6GHO;7F75; |
| rpoA BSU01430 | 1Z3E;3GFK;3IHQ;6WVJ;6WVK;6ZCA;6ZFB;7CKQ;7F75; |
| namA yqjM BSU23820 | 1Z41;1Z42;1Z44;1Z48; |
| yngHB BSU18239 | 1Z6H;1Z7T;2B8F;2B8G; |
| ohrR ykmA BSU13150 | 1Z91;1Z9C; |
| nfrA2 ycnD BSU03860 | 1ZCH; |
| ywlE BSU36930 ipc-31d | 1ZGG;4ETI;4ETN;4KK3;4KK4; |
| prsA BSU09950 | 1ZK6;4WO7; |
| racE glr murI BSU28390 | 1ZUW; |
| ccpA alsA amyR graR BSU29740 | 1ZVV;2FEP;3OQM;3OQN;3OQO; |
| yktB BSU14650 | 2A8E; |
| pyrB BSU15490 | 2AT2;3R7D;3R7F;3R7L; |
| yorR BSU20280 | 2AXP;3KB2; |
| codY BSU16170 | 2B0L;2B18;2GX5;2HGV;5LNH;5LOE;5LOJ;5LOO; |
| ribD ribG BSU23280 | 2B3Z;2D5N;3EX8;4G3M; |
| opuAC BSU03000 | 2B4L;2B4M;3CHG;5NXX; |
| ykuV BSU14230 | 2B5X;2B5Y; |
| ykuI BSU14090 | 2BAS;2W27; |
| yosS yojU BSU20020 | 2BAZ;2XX6;2XY3;2Y1T;4AO5; |
| hutI BSU39370 EE57B | 2BB0;2G3F; |
| yoaJ BSU18630 | 2BH0;3D30;4FER;4FFT;4FG2;4FG4; |
| ohrB ykzA yzzE BSU13160 | 2BJO; |
| nagB BSU35020 | 2BKV;2BKX; |
| rsbRA rsbR ycxR BSU04670 | 2BNL; |
| yukD BSU31900 | 2BPS; |
| bofC BSU27750 | 2BW2;7XT1; |
| rtpA yczA BSU02530 | 2BX9;2KO8;2ZP8;2ZP9; |
| citA BSU09440 | 2C6X; |
| lrpC ydaI BSU04250 | 2CFX; |
| mrgA BSU32990 | 2CHP; |
| walR yycF BSU40410 | 2D1V;2ZWM;3F6P; |
| yjcG BSU11850 | 2D4G; |
| uvrB dinA uvr BSU35170 | 2D7D;2NMV;3V4R; |
| yopT BSU20770 | 2DLB; |
| yqaI BSU26300 | 2DSM; |
| amyX BSU29930 | 2E8Y;2E8Z;2E9B; |
| yfmB BSU07530 | 2EUC; |
| yvdT BSU34480 | 2F07; |
| fapR ylpC BSU15880 | 2F3X;2F41; |
| trmB ytmQ BSU29900 | 2FCA;7NYB;7NZI;7NZJ; |
| perR ygaG BSU08730 | 2FE3;2RGV;3F8N; |
| mtnX ykrX BSU13600 | 2FEA; |
| ykuJ BSU14100 | 2FFG; |
| yycH BSU40390 | 2FGT; |
| acyP AcP yflL BSU07640 | 2FHM;2HLT;2HLU;3BR8; |
| yozE BSU19680 | 2FJ6; |
| hutU BSU39360 EE57A | 2FKN; |
| ywmB BSU36770 | 2FPN; |
| hpr catA scoC BSU09990 | 2FXA; |
| abh ylxT yzaA BSU14480 | 2FY9;2RO3; |
| dbpA deaD yxiN BSU39110 SS8E | 2G0C;2HJV;3MOJ; |
| yyaP BSU40760 | 2GD9; |
| yitF BSU10970 | 2GDQ;2GGE; |
| yqjZ BSU23680 | 2GO8; |
| yvfG BSU34210 | 2GSV;2JS1; |
| ypjD jojD BSU22500 | 2GTA; |
| tseB ypmB BSU22380 | 2GU3; |
| xkdM BSU12660 | 2GUJ;5LI2; |
| yybH BSU40640 | 2GXF; |
| trxA trx BSU28500 | 2GZY;2GZZ;2IPA;2VOC; |
| pucM yunM BSU32460 | 2H0E;2H0F;2H0J; |
| yonK BSU21060 | 2H4O; |
| yvyC yviH BSU35350 | 2HC5; |
| ynzC BSU17880 | 2HEP;2JVD;3BHP; |
| yorP BSU20300 | 2HEQ; |
| yppE BSU22270 | 2HFI;2IM8; |
| xkdW BSU12760 | 2HG7; |
| yjcQ BSU11950 | 2HGC; |
| der engA yphC BSU22840 | 2HJG;4DCS;4DCT;4DCU;4DCV;5M7H;5MBS;5X4B; |
| yqbF BSU26130 | 2HJQ; |
| ytcD BSU29030 | 2HZT; |
| yopX BSU20730 | 2I2L; |
| pdxK ywdB BSU38020 ipa-52r | 2I5B; |
| spoVG BSU00490 | 2IA9; |
| yckB BSU03380 | 2IEE; |
| yurK BSU32560 | 2IKK; |
| hemE BSU10120 | 2INF; |
| ykvR BSU13800 | 2JN9; |
| yxeF BSU39570 HS74F | 2JOZ; |
| yobA BSU18810 | 2JQO;4QY7; |
| tpx ytgI BSU29490 | 2JSY;2JSZ; |
| nfeD2 yuaF BSU31020 | 2K14; |
| srfAD srfA4 BSU03520 | 2K2Q;2RON; |
| yugI BSU31390 | 2K4K; |
| clpC mecB BSU00860 | 2K77;2Y1Q;2Y1R;3J3R;3J3S;3J3T;3J3U;3PXG;3PXI;5HBN;7ABR;8B3S;8OTK; |
| dnaI ORF311 ytxA BSU28980 | 2K7R;4M4W; |
| yndB BSU17730 | 2KEW;2KTE; |
| ylbL BSU15050 | 2KJP; |
| yutD BSU32310 | 2KL5; |
| rpoE BSU37160 | 2KRC;2M4K;4NC7;4NC8;6ZCA;6ZFB;7F75; |
| comA comA1 comAA BSU31680 | 2KRF;3ULQ; |
| pxpB kipI ycsJ BSU04080 | 2KWA;2ZP2; |
| ydhK BSU05790 | 2KY9;4FIB;4MDW; |
| msrB yppQ BSU21680 | 2KZN;3E0O; |
| spoIIID BSU36420 | 2L0K; |
| tatAd yczB BSU02630 | 2L16; |
| eglS bglC gld BSU18130 | 2L8A;3PZT;3PZU;3PZV;6UFV;6UFW; |
| yolA BSU21540 | 2LR4; |
| rpoB crsE rfm BSU01070 | 2LY7;6WVJ;6WVK;6ZCA;6ZFB;7CKQ;7F75; |
| yqzG BSU24650 | 2LYX; |
| ykzF BSU14120 | 2LZF; |
| minC BSU28000 | 2M4I; |
| sunA yolG BSU21480 | 2MIJ; |
| mciZ BSU_23616 | 2MRW;4U39; |
| nusA BSU16600 | 2MT4; |
| spoVM BSU15810 | 2MVH;2MVJ; |
| pfyP ytvA BSU30340 | 2MWG;2PR5;2PR6;4GCZ; |
| yqgQ BSU24860 | 2NN4; |
| pdxS yaaD BSU00110 | 2NV1;2NV2; |
| ytmB BSU30570 | 2NWA; |
| ypsA BSU22190 | 2NX2; |
| yokD BSU21630 | 2NYG; |
| csaA BSU19040 | 2NZH;2NZO; |
| yxiM BSU39120 SS8D | 2O14; |
| znuA adcA ycdH BSU02850 | 2O1E; |
| tcyK ytmK BSU29370 | 2O1M; |
| ybbH BSU01690 | 2O3F; |
| yycI BSU40380 | 2O3O; |
| fra ydhG BSU05750 | 2OC6; |
| yheA BSU09800 | 2OEE; |
| treR yfxA BSU07820 | 2OGG; |
| yueI BSU31770 | 2OHW; |
| cggR yvbQ BSU33950 | 2OKG;3BXE;3BXF;3BXG;3BXH;7OYK;8R3G; |
| mtnK ykrT BSU13560 | 2OLC;2PU8;2PUI;2PUL;2PUN;2PUP; |
| ywhB BSU37540 | 2OP8;2OPA; |
| yxeI BSU39540 LP9A | 2OQC; |
| padC pad yveH BSU34400 | 2P8G;4ALB; |
| aadK BSU26790 HIR78_15755 | 2PBE; |
| feuA BSU01630 | 2PHZ;2WHY;2WI8;2XUZ;2XV1; |
| ymcA BSU17020 | 2PIH;6PRH;6PRK; |
| ylbP BSU15100 | 2PR1; |
| yobK BSU18990 | 2PRV; |
| yerB yecC BSU06570 | 2PSB; |
| flhF BSU16400 | 2PX0;2PX3;3SYN; |
| yrrB BSU27490 | 2Q7F; |
| cotI ytaA BSU30920 | 2Q83; |
| yizA yucC BSU10800 | 2QE9; |
| ykqA ylxU yzaB BSU14500 | 2QIK; |
| nap BSU05440 | 2R11; |
| manP yjdD BSU12010 | 2R48; |
| fruA BSU14400 | 2R4Q; |
| sodA yqgD BSU25020 | 2RCV; |
| ftsZ BSU15290 | 2RHH;2RHJ;2RHL;2RHO;2VAM;2VXY;4U39; |
| dpaA spoVFA BSU16730 orfY | 2RIR; |
| spoVT yabL BSU00560 | 2RO5;2W1R;2W1T; |
| yoeB BSU18380 | 2RSX;8I2E;8I2F;8WT4; |
| ggt BSU18410 | 2V36;3A75;3WHQ;3WHR;3WHS; |
| dnaD BSU22350 | 2V79;8OJJ; |
| hisG BSU34920 | 2VD2; |
| nagA BSU35010 | 2VHL; |
| kinA gsiC scoD spoIIF spoIIJ BSU13990 | 2VLG; |
| rex ydiH BSU05970 | 2VT2;2VT3; |
| ltaS2 yflE BSU07710 | 2W8D; |
| gmuG ydhT BSU05880 | 2WHK;3CBW; |
| divIVA ylmJ BSU15420 | 2WUJ;2WUK; |
| nagR yvoA BSU35030 | 2WV0;4U0V;4U0W;4U0Y;4WWC; |
| menD BSU30820 | 2X7J; |
| abn2 J3A yxiA BSU39330 | 2X8F;2X8S;2X8T;4COT; |
| yncF BSU17660 | 2XCD;2XCE;4AOO;4AOZ;4APZ;4B0H; |
| purD BSU06530 | 2XCL;2XD4; |
| ydaE BSU04200 | 2Y0O; |
| mecA BSU11520 | 2Y1R;3J3R;3J3S;3J3T;3J3U;3JTP;3PXG;3PXI; |
| cymR yrzC BSU27520 | 2Y75; |
| phoD ycbS BSU02620 | 2YEQ; |
| mtnA ykrS BSU13550 | 2YRF;2YVK; |
| sspC BSU19950 | 2Z3X; |
| yesW BSU07050 | 2Z8R;2Z8S;2ZUX; |
| yesX BSU07060 | 2ZUY; |
| mtnW ykrW BSU13590 | 2ZVI; |
| hfq ymaH BSU17340 | 3AHU;3HSB; |
| ytbE BSU29050 | 3B3D; |
| mhqN ydfN BSU05480 | 3BEM; |
| queC ykvJ BSU13720 | 3BL5; |
| nagZ ybbD yzbA BSU01660 | 3BMX;3LK6;3NVD;4GYJ;4GYK; |
| hemL hemK BSU28120 | 3BS8; |
| yydK BSU40130 | 3BWG; |
| yetF BSU07140 | 3C6F; |
| xynD BSU18160 | 3C7E;3C7F;3C7G;3C7H;3C7O; |
| scoA yxjD BSU38990 N15K | 3CDK; |
| scoB yxjE BSU38980 N15L | 3CDK; |
| yjhA BSU12180 | 3CFU; |
| phoR BSU29100 | 3CWF; |
| ydhD BSU05710 | 3CZ8; |
| yflH BSU07680 | 3D0W; |
| yvgN BSU33400 | 3D3F;3F7J; |
| yjoA BSU12410 | 3DKA; |
| yhaX BSU09830 | 3DNP; |
| bioK bioA BSU30230 | 3DOD;3DRD;3DU4;6WNN; |
| dltA dae BSU38500 ipa-5r | 3E7W;3E7X; |
| yfnB BSU07330 | 3ED5;3I76; |
| panE apbA ylbQ BSU15110 | 3EGO; |
| desK yocF BSU19190 | 3EHF;3EHG;3EHH;3EHJ;3GIE;3GIF;3GIG;5IUJ;5IUK;5IUL;5IUM;7SSI;7SSJ; |
| bioI CYP107H BSU30190 | 3EJB;3EJD;3EJE; |
| cdoA yubC BSU31140 | 3EQE;4QM8;4QM9; |
| stoA spoIVH BSU13840 | 3ERW; |
| bdbD yvgV BSU33480 | 3EU3;3EU4;3GH9;3GHA; |
| frlB yurP BSU32610 | 3EUA; |
| xkdH BSU12620 | 3F3B; |
| ydeA BSU05110 | 3F5D; |
| lplD BSU07130 | 3FEF; |
| brxA yphP BSU21860 | 3FHK; |
| ccpN yqzB BSU25250 | 3FV6;3FWR;3FWS; |
| degV yviA BSU35480 | 3FYS; |
| fhuD BSU33320 | 3G9Q;3HXP; |
| nfi ywqL BSU36170 | 3GA2; |
| mutL BSU17050 | 3GAB;3KDG;3KDK; |
| iolW yvaA BSU33530 | 3GDO;3GFG; |
| yclQ BSU03830 | 3GFV; |
| xynC ynfF BSU18150 | 3GTN;3KL0;3KL3;3KL5; |
| yfkN BSU07840 | 3GVE; |
| glcT ykwA BSU13880 | 3GWH;3RIO;3UFE; |
| ntdB yhjK BSU10540 | 3GYG; |
| bacB ywfC BSU37730 ipa-81d | 3H7J;3H7Y;3H9A; |
| xlyA BSU12810 | 3HMB;3RDR; |
| pksH BSU17160 | 3HP0; |
| ypwA BSU22080 | 3HQ2; |
| cgoX hemG hemY BSU10140 | 3I6D; |
| pucG yurG BSU32520 | 3ISL; |
| ipi BSU11130 | 3ISY; |
| rplX BSU01270 | 3J3V;3J3W;3J9W;5NJT;6HA1;6HA8;6HTQ;6PPF;6PPK;6PVK;6TNN;6TPQ;7AQC;7AQD;7AS8;7AS9;7O5B;7OPE;7QV1;7QV2;7QV3;7S9U;7SAE;8BUU;8QCQ;8S1P;8S1U; |
| rplO BSU01350 | 3J3V;3J3W;3J9W;5NJT;6HA1;6HA8;6HTQ;6PPF;6PPK;6PVK;6TNN;6TPQ;7AQC;7AQD;7AS8;7AS9;7O5B;7OPE;7QV1;7QV2;7QV3;7S9U;7SAE;8BUU;8QCQ;8S1P;8S1U; |
| rplU BSU27960 | 3J3V;3J3W;3J9W;5NJT;6HA1;6HA8;6HTQ;6PPF;6PPK;6PVK;6TNN;6TPQ;7AQC;7AQD;7AS8;7AS9;7O5B;7OPE;7QV1;7QV2;7QV3;7S9U;7SAE;8BUU;8QCQ;8S1P;8S1U; |
| rplV BSU01210 | 3J3V;3J3W;3J9W;5NJT;6HA1;6HA8;6HTQ;6PPF;6PPK;6PVK;6TNN;6TPQ;7AQC;7AQD;7AS8;7AS9;7O5B;7OPE;7QV1;7QV2;7QV3;7S9U;7SAE;8BUU;8QCQ;8S1P;8S1U; |
| rplB BSU01190 | 3J3V;3J3W;3J9W;5NJT;6HA1;6HA8;6HTQ;6PPF;6PPK;6PVK;6TNN;6TPQ;7AQC;7AQD;7AS8;7AS9;7O5B;7OPE;7QV1;7QV2;7QV3;7S9U;7SAE;8BUU;8QCQ;8S1P;8S1U; |
| rplC BSU01160 | 3J3V;3J3W;3J9W;5NJT;6HA1;6HA8;6HTQ;6PPF;6PPK;6PVK;6TNN;6TPQ;7AQC;7AQD;7AS8;7AS9;7O5B;7OPE;7QV1;7QV2;7QV3;7S9U;7SAE;8BUU;8QCQ;8S1P;8S1U; |
| rplD BSU01170 | 3J3V;3J3W;3J9W;5NJT;6HA1;6HA8;6HTQ;6PPF;6PPK;6PVK;6TNN;6TPQ;7AQC;7AQD;7AS8;7AS9;7O5B;7OPE;7QV1;7QV2;7QV3;7S9U;7SAE;8BUU;8QCQ;8S1P;8S1U; |
| rplM BSU01490 | 3J3V;3J3W;3J9W;5NJT;6HA1;6HA8;6HTQ;6PPF;6PPK;6PVK;6TNN;6TPQ;7AQC;7AQD;7AS8;7AS9;7O5B;7OPE;7QV1;7QV2;7QV3;7S9U;7SAE;8BUU;8QCQ;8S1P;8S1U; |
| rplS BSU16040 | 3J3V;3J3W;3J9W;5NJT;6HA1;6HA8;6HTQ;6PPF;6PPK;6PVK;6TNN;6TPQ;7AQC;7AQD;7AS8;7AS9;7O5B;7OPE;7QV1;7QV2;7QV3;7S9U;7SAE;8BUU;8QCQ;8S1P;8S1U; |
| rpmF BSU15080 | 3J3V;3J3W;3J9W;5NJT;6HA1;6HA8;6HTQ;6PPF;6PPK;6PVK;6TNN;6TPQ;7AQC;7AQD;7AS8;7AS9;7O5B;7OPE;7QV1;7QV2;7QV3;7S9U;7SAE;8BUU;8QCQ;8S1P;8S1U; |
| rpmH BSU41060 | 3J3V;3J3W;3J9W;5NJT;6HA1;6HA8;6HTQ;6PPF;6PPK;6PVK;6TNN;6TPQ;7AQC;7AQD;7AS8;7AS9;7O5B;7OPE;7QV1;7QV2;7QV3;7S9U;7SAE;8BUU;8QCQ;8S1P;8S1U; |
| rpmC BSU01240 | 3J3V;3J3W;3J9W;5NJT;6HA1;6HA8;6HTQ;6PPF;6PPK;6PVK;6TNN;6TPQ;7AQC;7AQD;7AS8;7AS9;7O5B;7OPE;7QV1;7QV2;7QV3;7S9U;7SAE;8BUU;8QCQ;8S1P;8S1U; |
| rplN BSU01260 | 3J3V;3J3W;3J9W;5NJT;6HA1;6HA8;6HTQ;6PPF;6PPK;6PVK;6TNN;6TPQ;7AQC;7AQD;7AS8;7AS9;7O5B;7OPE;7QV1;7QV2;7QV3;7S9U;7SAE;8BUU;8QCQ;8S1P;8S1U; |
| rplQ BSU01440 | 3J3V;3J3W;3J9W;5NJT;6HA1;6HA8;6HTQ;6PPF;6PPK;6PVK;6TNN;6TPQ;7AQC;7AQD;7AS8;7AS9;7O5B;7OPE;7QV1;7QV2;7QV3;7S9U;7SAE;8BUU;8QCQ;8S1P;8S1U; |
| rplW BSU01180 | 3J3V;3J3W;3J9W;5NJT;6HA1;6HA8;6HTQ;6PPF;6PPK;6PVK;6TNN;6TPQ;7AQC;7AQD;7AS8;7AS9;7O5B;7OPE;7QV1;7QV2;7QV3;7S9U;7SAE;8BUU;8QCQ;8S1P;8S1U; |
| rplT BSU28850 | 3J3V;3J3W;3J9W;5NJT;6HA1;6HA8;6HTQ;6PPF;6PPK;6PVK;6TNN;6TPQ;7AQC;7AQD;7AS8;7AS9;7O5B;7OPE;7QV1;7QV2;7QV3;7S9U;7SAE;8BUU;8QCQ;8S1P;8S1U; |
| rplF BSU01310 | 3J3V;3J3W;3J9W;5NJT;6HA1;6HA8;6HTQ;6PPK;6TNN;6TPQ;7AQC;7AQD;7AS8;7AS9;7O5B;7OPE;7QV1;7QV2;7QV3;8BUU;8QCQ;8R55;8S1P;8S1U; |
| rplK relC tsp6 BSU01020 | 3J3V;3J3W;3J9W;7AQC;7AQD;7AS8;7AS9;7ASA;7O5B;7OPE; |
| rplA BSU01030 | 3J3V;3J3W;6HA8; |
| rpmD BSU01340 | 3J3V;3J9W;5NJT;6HA1;6HA8;6HTQ;6PPF;6PPK;6TNN;6TPQ;7AQC;7AQD;7AS8;7AS9;7O5B;7OPE;7QV1;7QV2;7QV3;7S9U;7SAE;8BUU;8QCQ;8S1P;8S1U; |
| rplR BSU01320 | 3J3V;3J9W;5NJT;6HA1;6HA8;6HTQ;6PPF;6PPK;6TNN;6TPQ;7AQC;7AQD;7AS8;7AS9;7O5B;7OPE;7QV1;7QV2;7QV3;8BUU;8QCQ;8S1P;8S1U; |
| rplE BSU01280 | 3J3V;3J9W;5NJT;6HA1;6HA8;6HTQ;6PPK;6TNN;6TPQ;7AQC;7AQD;7AS8;7AS9;7O5B;7OPE;7QV1;7QV2;7QV3;8BUU;8QCQ;8S1P;8S1U; |
| mifM yqzJ BSU23880 | 3J9W; |
| rpmA BSU27940 | 3J9W;5NJT;6HA1;6HA8;6HTQ;6PPF;6PPK;6TNN;6TPQ;7AQC;7AQD;7AS8;7AS9;7O5B;7OPE;7QV1;7QV2;7QV3;7S9U;7SAE;8BUU;8QCQ;8S1P;8S1U; |
| rplP BSU01230 | 3J9W;5NJT;6HA1;6HA8;6HTQ;6TNN;6TPQ;7AQC;7AQD;7AS8;7AS9;7O5B;7OPE;7QV1;7QV2;7QV3;8BUU;8QCQ;8S1P;8S1U; |
| rpmJ BSU01400 | 3J9W;5NJT;6HA1;6HA8;6HTQ;6TNN;6TPQ;7AQC;7AQD;7AS8;7AS9;7O5B;7OPE;7QV1;7QV2;7QV3;8BUU;8QCQ;8S1P;8S1U; |
| rpmGA rpmG1 BSU24900 | 3J9W;5NJT;6HA1;6HA8;6HTQ;6TNN;6TPQ;7AQC;7AQD;7AS8;7AS9;7O5B;7OPE;7QV1;7QV2;7QV3;8BUU;8QCQ;8S1P;8S1U; |
| rpmB yloT BSU15820 | 3J9W;5NJT;6HA1;6HA8;6HTQ;6TNN;6TPQ;7AQC;7AQD;7AS8;7AS9;7O5B;7OPE;7QV1;7QV2;7QV3;8BUU;8QCQ;8S1P;8S1U; |
| rpmI BSU28860 | 3J9W;5NJT;6HA1;6HA8;6HTQ;6TNN;6TPQ;7AQC;7AQD;7AS8;7AS9;7O5B;7OPE;7QV1;7QV2;7QV3;8BUU;8QCQ;8S1P;8S1U; |
| rpsD BSU29660 | 3J9W;5NJT;6HA1;6HA8;6HTQ;7O5B;7QV1;7QV2;7QV3;8BUU;8CDU;8CDV;8CEC;8CED;8CEE; |
| rpsP BSU15990 | 3J9W;5NJT;6HA1;6HA8;6HTQ;7O5B;7QV1;7QV2;7QV3;8BUU;8CDU;8CDV;8CEC;8CED;8CEE; |
| rpsL fun strA BSU01100 | 3J9W;5NJT;6HA1;6HA8;6HTQ;7O5B;7QV1;7QV2;7QV3;8BUU;8CDU;8CDV;8CEC;8CED;8CEE;8QCQ; |
| rpsE spcA BSU01330 | 3J9W;5NJT;6HA1;6HA8;6HTQ;7O5B;7QV1;7QV2;7QV3;8BUU;8CDU;8CDV;8CEC;8CED;8CEE;8QCQ; |
| rpsQ BSU01250 | 3J9W;5NJT;6HA1;6HA8;6HTQ;7O5B;7QV1;7QV2;7QV3;8BUU;8CDU;8CDV;8CEC;8CED;8CEE;8QCQ; |
| rpsH BSU01300 | 3J9W;5NJT;6HA1;6HA8;6HTQ;7O5B;7QV1;7QV2;7QV3;8BUU;8CDU;8CDV;8CEC;8CED;8CEE;8QCQ; |
| rpsM BSU01410 | 3J9W;5NJT;6HA1;6HA8;6HTQ;7O5B;7QV1;7QV2;7QV3;8BUU;8CDU;8CDV;8CEC;8CED;8CEE;8QCQ; |
| rpsF BSU40910 | 3J9W;5NJT;6HA1;6HA8;6HTQ;7O5B;7QV1;7QV2;7QV3;8BUU;8CDU;8CDV;8CEC;8CED;8CEE;8QCQ; |
| rpsG BSU01110 | 3J9W;5NJT;6HA1;6HA8;6HTQ;7O5B;7QV1;7QV2;7QV3;8BUU;8CDU;8CDV;8CEC;8CED;8CEE;8QCQ; |
| rpsI BSU01500 | 3J9W;5NJT;6HA1;6HA8;6HTQ;7O5B;7QV1;7QV2;7QV3;8BUU;8CDU;8CDV;8CEC;8CED;8CEE;8QCQ; |
| rpsJ tetA BSU01150 | 3J9W;5NJT;6HA1;6HA8;6HTQ;7O5B;7QV1;7QV2;7QV3;8BUU;8CDU;8CDV;8CEC;8CED;8CEE;8QCQ; |
| rpsO BSU16680 | 3J9W;5NJT;6HA1;6HA8;6HTQ;7O5B;7QV1;7QV2;7QV3;8BUU;8CDU;8CDV;8CEC;8CED;8CEE;8QCQ; |
| rpsR BSU40890 | 3J9W;5NJT;6HA1;6HA8;6HTQ;7O5B;7QV1;7QV2;7QV3;8BUU;8CDU;8CDV;8CEC;8CED;8CEE;8QCQ; |
| rpsS BSU01200 | 3J9W;5NJT;6HA1;6HA8;6HTQ;7O5B;7QV1;7QV2;7QV3;8BUU;8CDU;8CDV;8CEC;8CED;8CEE;8QCQ; |
| rpsT yqeO BSU25550 | 3J9W;5NJT;6HA1;6HA8;6HTQ;7O5B;7QV1;7QV2;7QV3;8BUU;8CDU;8CDV;8CEC;8CED;8CEE;8QCQ; |
| rpsK BSU01420 | 3J9W;5NJT;6HA1;6HA8;6HTQ;7O5B;7QV1;7QV2;7QV3;8BUU;8CDU;8CDV;8CEC;8CED;8CEE;8QCQ; |
| rpsN1 rpsN rpsNA rpsZ BSU01290 | 3J9W;5NJT;6HA1;6HA8;6HTQ;7O5B;7QV1;7QV2;7QV3;8BUU;8CDU;8CEC;8CED;8CEE;8QCQ; |
| rpsC BSU01220 | 3J9W;5NJT;6HA1;6HA8;6HTQ;7O5B;7QV1;7QV2;7QV3;8BUU;8CDU;8CEC;8CED;8CEE;8QCQ; |
| rpmE BSU37070 | 3J9W;5NJT;6HA1;6HA8;6HTQ;7O5B;7QV1;7QV2;7QV3;8QCQ;8S1P;8S1U; |
| rpsB BSU16490 | 3J9W;5NJT;6HA1;6HA8;6HTQ;7O5B;7QV2;8BUU;8CDU;8CEC;8CED;8CEE; |
| rplJ BSU01040 | 3J9W;5NJT;6TNN;6TPQ;7AS8;7AS9;7O5B;7OPE; |
| mecB ypbH BSU22970 | 3JTN;3JTO; |
| gudB ypcA BSU22960 | 3K8Z;7MFT; |
| rocG gudA yweB BSU37790 ipa-75d | 3K92; |
| yqbN BSU26040 | 3KLU; |
| clpP yvdN BSU34540 | 3KTG;3KTH;3KTI;3KTJ;3KTK;3TT6;3TT7;7FEP;7FEQ;7FER;7FES;7P80;7P81; |
| ymzC BSU17350 | 3KVP; |
| spoVAD BSU23410 | 3LM6; |
| thiN yloS BSU15800 | 3LM8; |
| yumC BSU32110 | 3LZW;3LZX; |
| hutG BSU39380 EE57C | 3M1R; |
| tagT ywtF BSU35840 | 3MEJ;4DE9;6MPS;6MPT;6UF5; |
| dacB BSU23190 | 3MFD; |
| flhA BSU16390 | 3MIX; |
| galM yoxA BSU18360 | 3MWX;4BZE;4BZF;4BZG;4BZH; |
| iolG idh BSU39700 E83G | 3MZ0;3NT2;3NT4;3NT5;3NTO;3NTQ;3NTR;4L8V;4L9R; |
| nfrA1 nfrA ywcG BSU38110 ipa-43d | 3N2S; |
| gerBC BSU35820 | 3N54; |
| yvdM BSU34550 | 3NAS; |
| cypX cyp134 cypB BSU35060 | 3NC3;3NC5;3NC6;3NC7; |
| yslB BSU28460 | 3NJC; |
| yxeA BSU39620 HS74A | 3NPP; |
| tagV yvhJ BSU35520 | 3NXH;6UF3; |
| bglS bgl licS BSU39070 N15B | 3O5S; |
| spoIISB BSU12820 | 3O6Q; |
| spoIISA ykaC BSU12830 | 3O6Q; |
| mhqO ydfO BSU05490 | 3OAJ; |
| fabL yfhR ygaA BSU08650 | 3OIC;3OID; |
| fabI yjbW BSU11720 | 3OIF;3OIG; |
| yutF BSU32290 | 3PDW; |
| opuCC yvbC BSU33810 | 3PPN;3PPO;3PPP;3PPQ;3PPR; |
| ldh lctE BSU03050 | 3PQD;3PQE;3PQF; |
| trpS BSU11420 | 3PRH; |
| rapH yeeH yzqA BSU06830 | 3Q15; |
| ywqE BSU36240 | 3QY6;3QY7; |
| opuBC proX BSU33710 | 3R6U;5NXY;6EYG;6EYH;6EYL;6EYQ; |
| tal orfU ywjH BSU37110 | 3R8R; |
| sppA yteI BSU29530 | 3RST;4KWB; |
| walK yycG BSU40400 | 3SL2; |
| ylxH BSU16410 | 3SYN; |
| spoIIE spoIIH BSU00640 | 3T91;3T9Q;5MQH;5UCG; |
| araR araC yvbS BSU33970 | 3TB6;4EGY;4EGZ;4H0E;5D4R;5D4S; |
| spoIIIAH BSU24360 | 3TUF;3UZ0; |
| spoIIQ ywnI BSU36550 | 3TUF;3UZ0; |
| addB BSU10620 | 3U44;3U4Q;4CEH;4CEI;4CEJ; |
| addA BSU10630 | 3U44;3U4Q;4CEH;4CEI;4CEJ; |
| bacG ywfH BSU37680 ipa-86r | 3U49;3U4C;3U4D; |
| rapF ywhJ BSU37460 | 3ULQ;4I9C;4I9E; |
| rulS rplGB ybaB ybxF BSU01090 | 3V7E;4LCK;4TZP;4TZV;4TZW;4TZZ; |
| rulQ rplGA ylxQ ymxC BSU16620 | 3V7Q; |
| ssbB ywpH BSU36310 | 3VDY; |
| bacD ywfE BSU37710 ipa-83d | 3VMM;3WNZ;3WO0;3WO1; |
| rsbX BSU04740 | 3W40;3W41;3W42;3W43;3W44;3W45; |
| yisP yucD BSU10810 | 3WE9; |
| smc ylqA BSU15940 | 3ZGX;5H66;5H67;5NMO;5NNV;5XG3; |
| scpA ypuG BSU23220 | 3ZGX;5H66;5H67;5XG3; |
| holA yqeN BSU25560 | 3ZH9; |
| sepF ylmF BSU15390 | 3ZIH;3ZII;8HZQ;8HZT; |
| ung ywdG BSU37970 ipa-57d | 3ZOQ;3ZOR; |
| rnjA ykqC BSU14530 | 3ZQ4; |
| eno BSU33900 | 4A3R;7XML; |
| pfkA pfk BSU29190 | 4A3S; |
| yybR BSU40540 | 4A5M;4A5N; |
| sacC BSU27030 | 4AZZ;4B1L;4B1M; |
| ymdB BSU16970 | 4B2O; |
| bslA yuaB BSU31080 | 4BHU; |
| dltC BSU38520 ipa-3r | 4BPF;4BPG;4BPH; |
| ctpB yvjB BSU35240 | 4C2C;4C2D;4C2E;4C2F;4C2G;4C2H; |
| ybfK BSU02260 | 4CCY; |
| deoD punB BSU19630 | 4D8V;4D8X;4D8Y;4D98;4D9H;4DA0;4DA6;4DA7;4DA8;4DAB;4DAE;4DAN;4DAO;4DAR; |
| gyrA cafB nalA BSU00070 | 4DDQ; |
| aldY yxkE BSU38830 | 4DNG; |
| nrdF BSU17390 | 4DR0; |
| yfkJ BSU07880 | 4ETM; |
| queF ykvM BSU13750 | 4F8B;4FGC;5UDG; |
| metQ yusA BSU32730 | 4GOT; |
| rapJ ycdE BSU02820 | 4GYO; |
| yncM BSU17690 | 4HFS; |
| rapI yddL BSU05010 | 4I1A; |
| pksJ pksK BSU17180 | 4J1Q;4J1S;4NA1;4NA2;4NA3;5KTK;8QK4; |
| ktrB yubG BSU31100 | 4J7C;5BUT;8K1S;8K1T;8K1U;8POO;8XMH;8XMI; |
| kinD ykvD BSU13660 | 4JGO;4JGP;4JGQ;4JGR; |
| ytkD mutTA BSU30630 | 4JZS;4JZT;4JZU;4JZV; |
| ntdA yhjL BSU10550 | 4K2B;4K2I;4K2M; |
| chaA yfkE BSU07920 | 4KJR;4KJS; |
| desR yocG BSU19200 | 4LDZ;4LE0;4LE1;4LE2;5IUJ;5IUK;5IUL;7SSI;7SSJ; |
| glnA BSU17460 | 4LNF;4LNI;4LNK;4LNN;4LNO;4S0R; |
| fabF yjaY BSU11340 | 4LS5;4LS6;4LS7;4LS8; |
| malL yvdL BSU34560 | 4M56;4M8U;4MAZ;4MB1;5WCZ;7LV6; |
| ndoAI mazE BSU04650 | 4ME7; |
| gabR ycnF BSU03890 | 4MGR;4N0B;4TV7;5T4J;5T4K;5X03;6UXZ; |
| nin comJ BSU03420 | 4MQD; |
| deoR yxxC BSU39430 | 4OQP;4OQQ;7BHY;8R7Y; |
| mrnC yazC BSU00950 | 4OUN;6TNN; |
| yodJ yokZ BSU19620 | 4OX3; |
| hmoB yhgC yixC BSU10100 | 4OZ5; |
| tgl yugV yuxF BSU31270 | 4P8I;4PA5; |
| yetJ BSU07200 | 4PGR;4PGS;4PGU;4PGV;4PGW;4TKQ;6NQ7;6NQ8;6NQ9; |
| pksI BSU17170 | 4Q1G;4Q1H;4Q1I;4Q1J;4Q1K; |
| nrgB BSU36520 | 4R25;4RX6; |
| glnR BSU17450 | 4R4E;7TFC; |
| ycdA BSU02780 | 4R4G; |
| melE msmE BSU30270 | 4R6H; |
| yesO BSU06970 | 4R6K;5Z6B;5Z6C; |
| darA yaaQ BSU00290 | 4RLE; |
| yjiB BSU12210 | 4RM4; |
| tnrA scgR BSU13310 | 4S0R; |
| sirA yneE yoxF BSU17900 | 4TPS; |
| dnaA dnaH BSU00010 | 4TPS;8BTG;8BV3; |
| dnaN dnaG BSU00020 | 4TR6;6E8D; |
| pksR BSU17220 | 4U3V; |
| ffh BSU15980 | 4UE4;7O5B;7O9F; |
| gpsB ypsB BSU22180 | 4UG3;5AN5;6GP7;6GPZ; |
| ezrA ytwP BSU29610 | 4UXV; |
| comFB comF2 BSU35460 | 4WAI; |
| bshC yllA BSU15120 | 4WBD; |
| yuiC BSU32070 | 4WJT;4WLI;4WLK; |
| yjgB BSU12150 | 4YGT; |
| pksG BSU17150 | 4YXQ;4YXT;4YXV; |
| pksS BSU17230 | 4YZR; |
| resE ypxE BSU23110 | 4ZR7; |
| menE BSU30790 | 5BUQ;5BUR;5BUS;5GTD;5X8F;5X8G; |
| exoA BSU40880 | 5CFE; |
| spo0M ygaI BSU08760 | 5CL2; |
| bshA jojH ypjH BSU22460 | 5D00;5D01; |
| queG ygaP yhbA BSU08910 | 5D08;5D0A;5D0B;5T8Y; |
| ispD yacM BSU00900 | 5DDT;5DDV;5HS2; |
| yjbM BSU11600 | 5DEC;5DED;5F2V; |
| yabA BSU00330 | 5DOL; |
| pksC BSU17100 | 5DZ6; |
| pksE BSU17120 | 5DZ7; |
| pksL outG pksA pksX BSU17190 | 5E1V;5E5N;5E6K;5ENY;5ERF; |
| ybxI ybdS BSU02090 | 5E2F;6W5E;6W5F; |
| ykoE BSU13230 | 5EDL; |
| pdp BSU39400 | 5EP8;5OLN; |
| yhjQ BSU10600 | 5FIG;6WKT; |
| bioW BSU30240 | 5FLG;5FLL;5FM0;5G1F; |
| cheR BSU22720 | 5FTW; |
| aroA BSU29750 | 5GMU;5GO2; |
| dnaG dnaE BSU25210 | 5GUJ; |
| yodB BSU19540 | 5HS7;5HS8;5HS9; |
| racA ywkC BSU37030 | 5I41;5I44; |
| nrnA ytqI BSU29250 | 5IPP;5IUF;5IZO;5J21; |
| alr2 yncD BSU17640 | 5IRP;6Q70;6Q71;6Q72; |
| bacC ywfD BSU37720 ipa-82d | 5ITV;5ITW; |
| ispF yacN BSU00910 | 5IWX;5IWY; |
| sufS csd yurW BSU32690 | 5J8Q;5XT5;5XT6;5ZS9;5ZSK;5ZSO;6KFY;6KFZ;7CEO;7CEP;7CEQ;7CER;7CES;7E6A;7E6B;7E6C;7E6D;7E6E;7E6F;7XEJ;7XEK;7XEL;7XEN;7XEP;7XET;7YB3; |
| rsiV yrhM BSU27130 | 5JEN; |
| pgpB yodM BSU19650 | 5JKI;6FMX; |
| mmgC yqiN BSU24150 | 5LNX; |
| mmgA yqiL BSU24170 | 5LP7; |
| hag BSU35360 | 5MAW;5WJT;6GOW; |
| bslB yweA BSU37800 ipa-74d | 5MKD; |
| yacP BSU00970 | 5MQ8;5MQ9; |
| fin subA yabK BSU00540 | 5MSL; |
| mmgE prpD yqiP BSU24130 | 5MUX; |
| tsaE ydiB BSU05910 | 5MVR;5NP9; |
| sigA rpoD BSU25200 | 5MWW;7CKQ;7F75; |
| csfB gin yaaM BSU00240 | 5N7Y; |
| yvyD hpf yviI BSU35310 orf189 | 5NJT; |
| spo0J BSU40960 | 5NOC;6SDK; |
| tasA cotN yqhF BSU24620 | 5OF1;5OF2;8AUR; |
| yndL BSU17820 | 5ONJ;5ONK;5ONL;6HRI;6HRJ; |
| sigW ybbL BSU01730 | 5OR5;5WUQ;5WUR;6JHE; |
| glpQ ybeD BSU02130 | 5T91;5T9B;5T9C; |
| queE ykvL BSU13740 | 5TGS;5TH5; |
| fusA fus BSU01120 | 5VH6; |
| ypfA jofA BSU22910 | 5VX6; |
| rsiW ybbM BSU01740 | 5WUQ;5WUR; |
| ycgT BSU03270 | 5XHU; |
| alsD BSU36000 | 5XNE; |
| ribT BSU23240 | 5XXR;5XXS; |
| tmcAL ylbM BSU15060 | 5Y0N;5Y0O;5Y0P;5Y0Q; |
| sirB ylnE BSU15620 | 5ZT7;5ZT8;5ZT9;5ZTA; |
| rok ykuW BSU14240 | 5ZUX;5ZUZ; |
| trmR yrrM BSU27360 | 5ZW3;5ZW4; |
| ctaA BSU14870 | 6A2J;6IED; |
| pucL yunL BSU32450 | 6A4M; |
| ssbA BSU40900 | 6BHW;6BHX; |
| spoIIIAB BSU24420 | 6BS9; |
| nrdE nrdA BSU17380 | 6CGL;6CGM;6CGN;6MT9;6MV9;6MVE;6MW3;6MYX; |
| spoIIIAF BSU24380 | 6DCS; |
| rocF BSU40320 | 6DKT;6NFP; |
| skfB ybcP ybcQ BSU01920 | 6EFN; |
| mgsA ypjF BSU22480 | 6F2C; |
| rny ymdA BSU16960 | 6F7T; |
| ywaC BSU38480 ipa-7d | 6FGK; |
| yabT BSU00660 | 6G4J; |
| gerM BSU28380 | 6GZ8;6GZB; |
| pdaC yjeA BSU12100 | 6H8L;6H8N; |
| vmlR expZ BSU05610 | 6HA8; |
| ywaD BSU38470 ipa-8r | 6HC6;6HC7; |
| yopK BSU20860 | 6HP3;6HP5;6HP7; |
| tapA yqhD yqxM BSU24640 | 6HQC;6QAY;8AIF; |
| relA BSU27600 | 6HTQ;6YXA;8ACU; |
| cdaA ybbP BSU01750 | 6HUW;7OJS;7OLH;8OGK;8OGM;8OGN;8OGO;8OGP;8OGQ;8OGR;8OGS;8OGT;8OGU;8OGV;8OGW;8OGY;8OGZ;8OH0;8OH1;8OHB;8OHC;8OHE;8OHF;8OHG;8OHH;8OHJ;8OHK;8OHL;8OHO; |
| xepA xkdY BSU12780 | 6I56;6IA5; |
| yomS BSU21240 | 6I5O; |
| ktrC ykqB ylxV yzaC BSU14510 | 6I8V; |
| bdhA ydjL BSU06240 | 6IE0; |
| rsmA ksgA BSU00420 | 6IFS;6IFT;6IFV;6IFW;7V2L;7V2M;7V2N;7V2O;7V2P;7V2Q; |
| rsbS ycxS BSU04680 | 6JHK; |
| efeB DyP efeN ywbN BSU38260 ipa-29d | 6KMM;6KMN;7DLK;7E5Q;7PKX;7PL0; |
| qoxB BSU38160 ipa-38d | 6KOB;6KOC;6KOE; |
| qoxA BSU38170 ipa-37d | 6KOB;6KOC;6KOE; |
| qoxC BSU38150 ipa-39d | 6KOB;6KOC;6KOE; |
| qoxD BSU38140 ipa-40d | 6KOB;6KOC;6KOE; |
| yjiC BSU12220 | 6KQW;6KQX;7BOV; |
| bacF ywfG BSU37690 ipa-85d | 6L1L;6L1N;6L1O; |
| rsbW BSU04720 | 6M36;6M37; |
| rsbV BSU04710 | 6M36;6M37; |
| yttP BSU29630 | 6MJ1; |
| bshB1 jojG ypjG BSU22470 | 6P2T;6ULL; |
| ylbF BSU14990 | 6PRK; |
| trmK yqfN BSU25180 | 6Q56; |
| patB BSU31440 | 6QP3; |
| gluP yqgP BSU24870 | 6R0J; |
| bmrA yvcC BSU34820 | 6R72;6R81;7BG4;7OW8;8CHB;8QOE; |
| kimA ydaO BSU04320 | 6S3K;8B70;8B71; |
| prfB BSU35290 | 6SZS; |
| yqkK BSU23540 | 6SZS; |
| mcsB yacI BSU00850 | 6TV6;8WTB;8WTC; |
| tagU lytR BSU35650 | 6UF6; |
| helD yvgS BSU33450 | 6VSX;6WVK;6ZCA;6ZFB; |
| penP BSU18800 | 6W2Z; |
| rpoC lpm std BSU01080 | 6WVJ;6WVK;6ZCA;6ZFB;7CKQ;7F75; |
| rpoZ yloH BSU15690 | 6WVJ;6WVK;7CKQ;7F75; |
| rpoY ykzG BSU14540 | 6WVK;6ZCA;6ZFB;7CKQ;7F75; |
| noc yyaA BSU40990 | 6Y93;7OL9; |
| msmX yxkG BSU38810 | 6YIR; |
| motB BSU13680 | 6YSL; |
| motA BSU13690 | 6YSL; |
| yukC BSU31890 | 6Z0F; |
| yprA BSU22220 | 6ZNP;6ZNQ;6ZNS; |
| khtT yhaT BSU09860 | 7AGV;7AGW;7AGY;7AHM;7AHT; |
| rqcH yloA BSU15640 | 7AQC;7AQD;7AS8;7AS9;7ASA;7OPE;8S1U; |
| rqcP yabO BSU00590 | 7AQC;7AS8;7ASA;7OPE; |
| yngB BSU18180 | 7B1R;7O2N; |
| yxeQ BSU39460 LP9I | 7BRA;7E9D; |
| hxlR BSU03470 | 7BZD;7BZE;7BZG; |
| csn BSU26890 | 7C6C;7C6D; |
| padR yfiO BSU08340 | 7CBV; |
| tadA yaaJ BSU00180 | 7CPH; |
| cssR BSU33010 | 7CX5; |
| tagH BSU35700 | 7DD0; |
| uppS yluA BSU16530 | 7JLI;7JLJ;7JLM;7JLR; |
| yheH BSU09720 | 7M33;8FHK;8FMV;8FPF;8SZC;8T1P;8T3K; |
| yheI BSU09710 | 7M33;8FHK;8FMV;8FPF;8SZC;8T1P;8T3K; |
| ycnI BSU03940 | 7ME6;7MEK;8UM6; |
| gltA BSU18450 | 7MFT; |
| gltB BSU18440 | 7MFT; |
| sunS yolJ BSU21450 | 7MSN;7MSP; |
| guaB gnaB BSU00090 | 7OJ1;7OJ2; |
| glmM ybbT BSU01770 | 7OJR;7OJS;7OLH;7OML; |
| yhcN BSU09150 | 7PEG; |
| pabB pab BSU00740 | 7PI1; |
| rlmP ysgA BSU28650 | 7QIU; |
| rplI BSU40500 | 7QV1;7QV3; |
| rpsU yqeX BSU25410 | 7QV1;7QV3; |
| mutSB mutS2 rqcU yshD BSU28580 | 7QV3;8R55; |
| ccrZ ytmP BSU29920 | 7S3L; |
| immR ydcN BSU04820 | 7T8I; |
| bceA barC ytsC BSU30380 | 7TCG;7TCH;8G3A;8G3B;8G3F;8G3L;8G4C;8G4D; |
| bceB barD ytsD BSU30370 | 7TCG;7TCH;8G3A;8G3B;8G3F;8G3L;8G4C;8G4D; |
| yojK BSU19420 | 7VM0;8H5D; |
| yjoB BSU12420 | 7W42;7W43;7W46; |
| yetL BSU07220 | 7WZE; |
| yopR BSU20790 | 8A0A;8BJ6;8BJV;8BPZ; |
| yydG BSU40170 | 8AI1;8AI2;8AI3;8AI4;8AI5;8AI6; |
| yydF BSU40180 | 8AI2;8AI4;8AI5; |
| phrE BSU25840 | 8ARE; |
| oppA spo0KA BSU11430 | 8ARE;8ARN; |
| pksD BSU17110 | 8AVZ;8AW0; |
| dppE dciAE BSU12960 | 8AY0;8AZB; |
| yjbA BSU11410 | 8B3S; |
| sigE spoIIGB BSU15320 | 8B3Z; |
| rnr vacB yvaJ BSU33610 | 8CDU;8CDV;8CEC;8CED;8CEE; |
| comEA comE1 BSU25590 | 8DFK; |
| lon2 lonB ysxF BSU28210 | 8DVH; |
| nusG BSU01010 | 8EHQ;8EJ3;8EOE;8EOF;8EXY; |
| bceS barB ytsB BSU30390 | 8G3A;8G3B;8G3F;8G3L;8G4C;8G4D; |
| mnmM ytqB BSU30490 | 8H0S;8H0T; |
| lytE cwlF papQ BSU09420 | 8I2D;8I2E;8I2F; |
| ytoQ BSU29850 | 8OF6; |
| rqcP2 ylmH BSU15410 | 8S1P;8S1U; |
| yisK BSU10750 | 8SKY;8SUT;8SUU; |
| rsgI ykrI BSU13460 | 8T9N; |
| yprB BSU22210 | 8UN9; |
| dfrA BSU21810 | 8UVZ; |
| fabHA fabH fabH1 yjaX BSU11330 | 8VD9;8VDA; |
| fabHB fabH2 yhfB BSU10170 | 8VDB; |
| spoIVFB bofB BSU27970 | 8VJL;8VJM; |
| sigK cisB spoIIIC spoIVCB BSU25760/BSU26390 | 8VJM; |
| gdh BSU03930 | 8W0N; |
| cspR ygaR BSU08930 | 8W9U; |
| cwlO yvcE yzkA BSU34800 | 8WT3;8WT4; |
| mcsA yacH BSU00840 | 8WTB;8WTC; |
| gapB BSU29020 | 8WWZ; |
| yrpD BSU26820 | 8XAI; |

**Supplementary Table 3.** GPR-PDB structure associations: The list of GPR Associations, PDB IDs and its corresponding DOI.

| GPR Association | PDB ID | DOI of Structure Articles |
| --- | --- | --- |
| BSU06170 or BSU05860 | &1XC3;3LM9;3OHR; |  |
| BSU06230 or BSU39710 |  |  |
| BSU06370 or BSU29990 |  |  |
| BSU06370 or BSU29990 |  |  |
| BSU07570 or BSU04470 |  |  |
| BSU07570 or BSU04470 |  |  |
| BSU07570 or BSU18540 |  |  |
| BSU07610 or BSU39060 |  |  |
| BSU09130 or BSU10220 |  |  |
| BSU09410 or BSU05740 |  |  |
| BSU09410 or BSU05740 |  |  |
| BSU09440 or BSU29140 | 2C6X;& |  |
| BSU09530 or BSU39780 | 1PZ1;&1PYF;1PZ0; |  |
| BSU09530 or BSU39780 | 1PZ1;&1PYF;1PZ0; | <https://doi.org/10.1016/j.jmb.2004.01.059> |
| BSU10220 or BSU02340 |  |  |
| BSU11710 or BSU38020 | &2I5B; |  |
| BSU12020 or BSU35790 | &1QWR; |  |
| BSU13120 or BSU18470 |  |  |
| BSU13850 or BSU33490 |  |  |
| BSU14880 or BSU12080 |  |  |
| BSU15480 or BSU36470 |  |  |
| BSU17660 or BSU20020 | 2XCD;2XCE;4AOO;4AOZ;4APZ;4B0H;&2BAZ;2XX6;2XY3;2Y1T;4AO5; | https://doi.org/10.1107/S0907444910026272 https://doi.org/10.1107/S090744491300735X <https://doi.org/10.1107/S0907444911003234> |
| BSU17680 or BSU21820 | 1B02;1BKO;1BKP;1BSF;1BSP;& |  |
| BSU18220 or BSU28540 |  |  |
| BSU18220 or BSU28540 |  |  |
| BSU18220 or BSU28540 |  |  |
| BSU18220 or BSU28540 |  |  |
| BSU18220 or BSU28540 |  |  |
| BSU18220 or BSU28540 |  |  |
| BSU18220 or BSU28540 |  |  |
| BSU18800 or BSU02090 | 6W2Z;&5E2F;6W5E;6W5F; | <https://doi.org/10.1016/j.jsb.2020.107544> |
| BSU19630 or BSU23490 | 4D8V;4D8X;4D8Y;4D98;4D9H;4DA0;4DA6;4DA7;4DA8;4DAB;4DAE;4DAN;4DAO;4DAR;& |  |
| BSU19630 or BSU23490 | 4D8V;4D8X;4D8Y;4D98;4D9H;4DA0;4DA6;4DA7;4DA8;4DAB;4DAE;4DAN;4DAO;4DAR;& |  |
| BSU19630 or BSU23490 | 4D8V;4D8X;4D8Y;4D98;4D9H;4DA0;4DA6;4DA7;4DA8;4DAB;4DAE;4DAN;4DAO;4DAR;& |  |
| BSU22620 or BSU34890 |  |  |
| BSU22620 or BSU34890 |  |  |
| BSU22890 or BSU16510 |  |  |
| BSU22890 or BSU16510 |  |  |
| BSU22960 or BSU37790 | 3K8Z;7MFT;&3K92; | <https://doi.org/10.1016/j.jmb.2010.05.055> <https://doi.org/10.1038/s41589-021-00919-y> |
| BSU23490 or BSU19630 | &4D8V;4D8X;4D8Y;4D98;4D9H;4DA0;4DA6;4DA7;4DA8;4DAB;4DAE;4DAN;4DAO;4DAR; |  |
| BSU23490 or BSU19630 | &4D8V;4D8X;4D8Y;4D98;4D9H;4DA0;4DA6;4DA7;4DA8;4DAB;4DAE;4DAN;4DAO;4DAR; |  |
| BSU23490 or BSU19630 | &4D8V;4D8X;4D8Y;4D98;4D9H;4DA0;4DA6;4DA7;4DA8;4DAB;4DAE;4DAN;4DAO;4DAR; |  |
| BSU23490 or BSU19630 | &4D8V;4D8X;4D8Y;4D98;4D9H;4DA0;4DA6;4DA7;4DA8;4DAB;4DAE;4DAN;4DAO;4DAR; |  |
| BSU23490 or BSU19630 | &4D8V;4D8X;4D8Y;4D98;4D9H;4DA0;4DA6;4DA7;4DA8;4DAB;4DAE;4DAN;4DAO;4DAR; |  |
| BSU23490 or BSU19630 | &4D8V;4D8X;4D8Y;4D98;4D9H;4DA0;4DA6;4DA7;4DA8;4DAB;4DAE;4DAN;4DAO;4DAR; |  |
| BSU23920 or BSU18210 |  |  |
| BSU24090 or BSU37660 | &1TD9;1XCO; |  |
| BSU24150 or BSU37170 | 5LNX;& |  |
| BSU24150 or BSU37170 | 5LNX;& |  |
| BSU24150 or BSU37170 | 5LNX;& |  |
| BSU25500 or BSU09840 |  |  |
| BSU26810 or BSU28390 | &1ZUW; |  |
| BSU26970 or BSU27010 |  |  |
| BSU28120 or BSU08710 | 3BS8;& |  |
| BSU28540 or BSU18220 |  |  |
| BSU28540 or BSU18220 |  |  |
| BSU28720 or BSU28510 |  |  |
| BSU29050 or BSU33400 | 3B3D;&3D3F;3F7J; | <https://doi.org/10.1002/pro.178> |
| BSU29300 or BSU16670 |  |  |
| BSU30830 or BSU31990 |  |  |
| BSU30870 or BSU38860 |  |  |
| BSU30970 or BSU30960 |  |  |
| BSU32440 or BSU32430 |  |  |
| BSU32450 or BSU32460 | 6A4M;&2H0E;2H0F;2H0J; | <https://doi.org/10.1134/S1063774519070149> <https://doi.org/10.1073/pnas.0600523103> |
| BSU32840 or BSU24160 |  |  |
| BSU33240 or BSU18670 | 1J58;1L3J;1UW8;2UY8;2UY9;2UYA;2UYB;2V09;3S0M;4MET;5HI0;5VG3;6TZP;6UFI;& |  |
| BSU33430 or BSU33440 |  |  |
| BSU33500 or BSU03950 | 1JWW;1KQK;1OPZ;1OQ3;1OQ6;1P6T;2RML;2VOY;& |  |
| BSU34890 or BSU22620 |  |  |
| BSU36120 or BSU36130 |  |  |
| BSU36760 or BSU37100 |  |  |
| BSU37120 or BSU39670 |  |  |
| BSU37170 or BSU24150 | &5LNX; |  |
| BSU37170 or BSU24150 | &5LNX; |  |
| BSU37170 or BSU24150 | &5LNX; |  |
| BSU37320 or BSU38060 |  |  |
| BSU38020 or BSU11710 | 2I5B;& |  |
| BSU38550 or BSU02390 |  |  |
| BSU38550 or BSU02390 |  |  |
| BSU38560 or BSU40110 |  |  |
| BSU38560 or BSU40110 |  |  |
| BSU38560 or BSU40110 |  |  |
| BSU38770 or BSU26860 |  |  |
| BSU39390 or BSU02400 |  |  |
| BSU39670 or BSU37120 |  |  |
| BSU40070 or BSU19520 |  |  |
| BSU00110 and BSU00120 | 2NV1;2NV2;&1R9G;2NV0;2NV2; | https://doi.org/10.1073/pnas.0604950103 https://doi.org/10.1074/jbc.M310311200 |
| BSU00740 and BSU00750 | 7PI1;& |  |
| BSU00750 and BSU22680 |  |  |
| BSU02750 and BSU02760 |  |  |
| BSU03300 and BSU03290 |  |  |
| BSU06420 and BSU06430 |  |  |
| BSU15530 and BSU15540 |  |  |
| BSU15850 and BSU15860 |  |  |
| BSU16090 and BSU16100 |  |  |
| BSU16730 and BSU16740 | 2RIR;& |  |
| BSU18450 and BSU18440 | 7MFT;&7MFT; |  |
| BSU22640 and BSU22630 |  |  |
| BSU22760 and BSU22740 |  |  |
| BSU23270 and BSU23250 | &1RVV;1ZIS; |  |
| BSU23270 and BSU23250 | &1RVV;1ZIS; |  |
| BSU28260 and BSU28250 |  |  |
| BSU28260 and BSU28250 |  |  |
| BSU35690 and BSU35680 |  |  |
| BSU38990 and BSU38980 | 3CDK;&3CDK; |  |
| BSU01580 or BSU15580 or BSU34660 |  |  |
| BSU04470 or BSU38770 or BSU07570 |  |  |
| BSU04520 or BSU18260 or BSU32820 |  |  |
| BSU08820 or BSU39050 or BSU38630 |  |  |
| BSU12910 or BSU18480 or BSU23800 |  |  |
| BSU13300 or BSU08000 or BSU24740 |  |  |
| BSU16760 or BSU28470 or BSU03790 |  |  |
| BSU18260 or BSU32820 or BSU04520 |  |  |
| BSU19310 or BSU38830 or BSU39860 | &4DNG; |  |
| BSU19310 or BSU38830 or BSU39860 | &4DNG;&&4DNG; |  |
| BSU19310 or BSU39860 or BSU38830 | &&4DNG; |  |
| BSU22690 or BSU27910 or BSU29750 | 1COM;1DBF;1FNJ;1FNK;2CHS;2CHT;3ZO8;3ZOP;3ZP4;3ZP7;&&5GMU;5GO2; | https://doi.org/10.1006/jmbi.1994.1462 https://doi.org/10.1107/s0907444900004625 https://doi.org/10.1074/jbc.M006351200 <https://doi.org/10.1073/pnas.90.18.8600> https://doi.org/10.1073/pnas.1408512111 <https://doi.org/10.1038/s41598-017-06578-1> |
| BSU27810 or BSU02420 or BSU18120 |  |  |
| BSU27810 or BSU02420 or BSU18120 |  |  |
| BSU30540 or BSU10790 or BSU39920 |  |  |
| BSU32200 or BSU12290 or BSU32100 |  |  |
| BSU33330 or BSU40330 or BSU37760 |  |  |
| BSU34790 or BSU32110 or BSU03270 | &3LZW;3LZX;&5XHU; | <https://doi.org/10.1002/pro.508> |
| BSU36030 or BSU05340 or BSU25790 |  |  |
| BSU37190 or BSU37240 or BSU36590 |  |  |
| BSU38110 or BSU03860 or BSU07830 | 3N2S;&1ZCH;& | https://doi.org/10.1016/j.febslet.2010.08.019 https://doi.org/10.1021/bi0510835 |
| BSU38830 or BSU19310 or BSU39860 | 4DNG;&& |  |
| BSU38830 or BSU39860 or BSU19310 | 4DNG;&& |  |
| BSU38830 or BSU39860 or BSU19310 | 4DNG;&& |  |
| BSU38830 or BSU39860 or BSU19310 | 4DNG;&& |  |
| BSU39020 or BSU39410 or BSU32180 |  |  |
| BSU39020 or BSU39410 or BSU32180 |  |  |
| BSU39410 or BSU32180 or BSU39020 |  |  |
| BSU39410 or BSU39020 or BSU32180 |  |  |
| BSU39860 or BSU19310 or BSU38830 | &&4DNG; |  |
| BSU39860 or BSU19310 or BSU38830 | &&4DNG; |  |
| BSU39860 or BSU19310 or BSU38830 | &&4DNG; |  |
| BSU39860 or BSU19310 or BSU38830 | &&4DNG; |  |
| BSU39860 or BSU38830 or BSU19310 | &4DNG;& |  |
| BSU39860 or BSU38830 or BSU19310 | &4DNG;& |  |
| BSU39860 or BSU38830 or BSU19310 | &4DNG;& |  |
| BSU39860 or BSU38830 or BSU19310 | &4DNG;& |  |
| BSU39860 or BSU38830 or BSU19310 | &4DNG;& |  |
| BSU39860 or BSU38830 or BSU19310 | &4DNG;& |  |
| BSU39860 or BSU38830 or BSU19310 | &4DNG;& |  |
| BSU39860 or BSU38830 or BSU19310 | &4DNG;& |  |
| BSU39860 or BSU38830 or BSU19310 | &4DNG;& |  |
| BSU39860 or BSU38830 or BSU19310 | &4DNG;& |  |
| BSU39860 or BSU38830 or BSU19310 | &4DNG;& |  |
| BSU39860 or BSU38830 or BSU19310 | &4DNG;& |  |
| BSU39860 or BSU38830 or BSU19310 | &4DNG;& |  |
| BSU39860 or BSU38830 or BSU19310 | &4DNG;& |  |
| BSU39860 or BSU38830 or BSU19310 | &4DNG;& |  |
| BSU39860 or BSU38830 or BSU19310 | &4DNG;& |  |
| BSU01630 and BSU01620 and BSU01610 | 2PHZ;2WHY;2WI8;2XUZ;2XV1;&& |  |
| BSU02350 and BSU13900 and BSU13910 | &1JEM;1KKL;1KKM;1SPH;2FEP;2HID;2HPR;3OQM;3OQN;3OQO;& |  |
| BSU02850 and BSU02860 and BSU02870 | 2O1E;&& |  |
| BSU03981 and BSU13900 and BSU13910 | &1JEM;1KKL;1KKM;1SPH;2FEP;2HID;2HPR;3OQM;3OQN;3OQO;& |  |
| BSU06480 and BSU06470 and BSU06460 | &&1T4A;1TWJ; |  |
| BSU06970 and BSU06980 and BSU06990 | 4R6K;5Z6B;5Z6C;&& |  |
| BSU07700 and BSU13900 and BSU13910 | &1JEM;1KKL;1KKM;1SPH;2FEP;2HID;2HPR;3OQM;3OQN;3OQO;& |  |
| BSU08200 and BSU13900 and BSU13910 | &1JEM;1KKL;1KKM;1SPH;2FEP;2HID;2HPR;3OQM;3OQN;3OQO;& |  |
| BSU08460 and BSU08440 and BSU08450 |  |  |
| BSU08840 and BSU08830 and BSU08850 |  |  |
| BSU08840 and BSU08830 and BSU08850 |  |  |
| BSU08840 and BSU08830 and BSU08850 |  |  |
| BSU08840 and BSU08830 and BSU08850 |  |  |
| BSU08840 and BSU08830 and BSU08850 |  |  |
| BSU08840 and BSU08830 and BSU08850 |  |  |
| BSU08840 and BSU08830 and BSU08850 |  |  |
| BSU08840 and BSU08830 and BSU08850 |  |  |
| BSU08840 and BSU08830 and BSU08850 |  |  |
| BSU12010 and BSU13900 and BSU13910 | 2R48;&1JEM;1KKL;1KKM;1SPH;2FEP;2HID;2HPR;3OQM;3OQN;3OQO;& |  |
| BSU13890 and BSU13900 and BSU13910 | 1AX3;1GPR;&1JEM;1KKL;1KKM;1SPH;2FEP;2HID;2HPR;3OQM;3OQN;3OQO;& | https://doi.org/10.1002/(SICI)1097-0134(19980515)31:3<258::AID-PROT3>3.0.CO;2-F https://doi.org/10.1016/S0021-9258(18)35837-X https://doi.org/10.1002/pro.5560061006 https://doi.org/10.1073/pnas.192368699 https://doi.org/10.1016/s0969-2126(94)00122-7 https://doi.org/10.1002/prot.21001 https://doi.org/10.1093/nar/gkq1177 |
| BSU13900 and BSU13910 and BSU07800 | 1JEM;1KKL;1KKM;1SPH;2FEP;2HID;2HPR;3OQM;3OQN;3OQO;&& |  |
| BSU17380 and BSU17390 and BSU17370 | 6CGL;6CGM;6CGN;6MT9;6MV9;6MVE;6MW3;6MYX;&4DR0;&1RLJ; | https://doi.org/10.1073/pnas.1800356115 https://doi.org/10.1038/s41467-019-10568-4 https://doi.org/10.1021/bi201925t |
| BSU17380 and BSU17390 and BSU17370 | 6CGL;6CGM;6CGN;6MT9;6MV9;6MVE;6MW3;6MYX;&4DR0;&1RLJ; | https://doi.org/10.1073/pnas.1800356115 https://doi.org/10.1038/s41467-019-10568-4 https://doi.org/10.1021/bi201925t |
| BSU17380 and BSU17390 and BSU17370 | 6CGL;6CGM;6CGN;6MT9;6MV9;6MVE;6MW3;6MYX;&4DR0;&1RLJ; | https://doi.org/10.1073/pnas.1800356115 https://doi.org/10.1038/s41467-019-10568-4 https://doi.org/10.1021/bi201925t |
| BSU17380 and BSU17390 and BSU17370 | 6CGL;6CGM;6CGN;6MT9;6MV9;6MVE;6MW3;6MYX;&4DR0;&1RLJ; | https://doi.org/10.1073/pnas.1800356115 https://doi.org/10.1038/s41467-019-10568-4 https://doi.org/10.1021/bi201925t |
| BSU19370 and BSU19360 and BSU14610 |  |  |
| BSU22560 and BSU22550 and BSU22540 |  |  |
| BSU23980 and BSU23970 and BSU23960 |  |  |
| BSU28450 and BSU28440 and BSU28430 |  |  |
| BSU28450 and BSU28440 and BSU28430 |  |  |
| BSU28750 and BSU28740 and BSU28730 |  |  |
| BSU33160 and BSU33170 and BSU33180 |  |  |
| BSU33370 and BSU33380 and BSU33390 |  |  |
| BSU36660 and BSU36650 and BSU36640 |  |  |
| BSU38280 and BSU38270 and BSU38260 | &&6KMM;6KMN;7DLK;7E5Q;7PKX;7PL0; |  |
| BSU39270 and BSU13900 and BSU13910 | &1JEM;1KKL;1KKM;1SPH;2FEP;2HID;2HPR;3OQM;3OQN;3OQO;& |  |
| BSU39270 and BSU13900 and BSU13910 | &1JEM;1KKL;1KKM;1SPH;2FEP;2HID;2HPR;3OQM;3OQN;3OQO;& |  |
| BSU06660 or BSU30530 or BSU03220 or BSU19580 |  |  |
| BSU10270 or BSU10360 or BSU18250 or BSU28560 |  |  |
| BSU10270 or BSU10360 or BSU18250 or BSU28560 |  |  |
| BSU10270 or BSU10360 or BSU18250 or BSU28560 |  |  |
| BSU10270 or BSU10360 or BSU18250 or BSU28560 |  |  |
| BSU10270 or BSU10360 or BSU18250 or BSU28560 |  |  |
| BSU10360 or BSU10270 or BSU18250 or BSU28560 |  |  |
| BSU10360 or BSU18250 or BSU10270 or BSU28560 |  |  |
| BSU10360 or BSU18250 or BSU28560 or BSU10270 |  |  |
| BSU18250 or BSU10270 or BSU10360 or BSU28560 |  |  |
| BSU18250 or BSU10360 or BSU28560 or BSU10270 |  |  |
| BSU28560 or BSU10270 or BSU10360 or BSU18250 |  |  |
| BSU28560 or BSU10270 or BSU10360 or BSU18250 |  |  |
| BSU28560 or BSU18250 or BSU10270 or BSU10360 |  |  |
| BSU28560 or BSU18250 or BSU10360 or BSU10270 |  |  |
| BSU12930 and BSU12940 and BSU12950 and BSU12960 | &&&8AY0;8AZB; |  |
| BSU12930 and BSU12940 and BSU12950 and BSU12960 | &&&8AY0;8AZB; |  |
| BSU12930 and BSU12940 and BSU12950 and BSU12960 | &&&8AY0;8AZB; |  |
| BSU12930 and BSU12940 and BSU12950 and BSU12960 | &&&8AY0;8AZB; |  |
| BSU12930 and BSU12940 and BSU12950 and BSU12960 | &&&8AY0;8AZB; |  |
| BSU12930 and BSU12940 and BSU12950 and BSU12960 | &&&8AY0;8AZB; |  |
| BSU12930 and BSU12940 and BSU12950 and BSU12960 | &&&8AY0;8AZB; |  |
| BSU12930 and BSU12940 and BSU12950 and BSU12960 | &&&8AY0;8AZB; |  |
| BSU12930 and BSU12940 and BSU12950 and BSU12960 | &&&8AY0;8AZB; |  |
| BSU12930 and BSU12940 and BSU12950 and BSU12960 | &&&8AY0;8AZB; |  |
| BSU12930 and BSU12940 and BSU12950 and BSU12960 | &&&8AY0;8AZB; |  |
| BSU12930 and BSU12940 and BSU12950 and BSU12960 | &&&8AY0;8AZB; |  |
| BSU12930 and BSU12940 and BSU12950 and BSU12960 | &&&8AY0;8AZB; |  |
| BSU12930 and BSU12940 and BSU12950 and BSU12960 | &&&8AY0;8AZB; |  |
| BSU12930 and BSU12940 and BSU12950 and BSU12960 | &&&8AY0;8AZB; |  |
| BSU14890 and BSU14900 and BSU14910 and BSU14920 |  |  |
| BSU24050 and BSU24040 and BSU24030 and BSU24060 |  |  |
| BSU24050 and BSU24040 and BSU24030 and BSU24060 |  |  |
| BSU24050 and BSU24040 and BSU24030 and BSU24060 |  |  |
| BSU30770 and BSU30760 and BSU30750 and BSU30740 |  |  |
| BSU33310 and BSU33290 and BSU33300 and BSU39610 |  |  |
| BSU33830 and BSU33820 and BSU33810 and BSU33800 | &&3PPN;3PPO;3PPP;3PPQ;3PPR;& |  |
| BSU33830 and BSU33820 and BSU33810 and BSU33800 | &&3PPN;3PPO;3PPP;3PPQ;3PPR;& |  |
| BSU33830 and BSU33820 and BSU33810 and BSU33800 | &&3PPN;3PPO;3PPP;3PPQ;3PPR;& |  |
| BSU33830 and BSU33820 and BSU33810 and BSU33800 | &&3PPN;3PPO;3PPP;3PPQ;3PPR;& |  |
| BSU33830 and BSU33820 and BSU33810 and BSU33800 | &&3PPN;3PPO;3PPP;3PPQ;3PPR;& |  |
| BSU35940 and BSU35960 and BSU35950 and BSU35930 | &&&1OGC;1OGD;1OGE;1OGF; |  |
| BSU35940 and BSU35960 and BSU35950 and BSU35930 | &&&1OGC;1OGD;1OGE;1OGF; |  |
| BSU35940 and BSU35960 and BSU35950 and BSU35930 | &&&1OGC;1OGD;1OGE;1OGF; |  |
| BSU38170 and BSU38160 and BSU38150 and BSU38140 | 6KOB;6KOC;6KOE;&6KOB;6KOC;6KOE;&6KOB;6KOC;6KOE;&6KOB;6KOC;6KOE; | <https://doi.org/10.1073/pnas.1915013117> |
| BSU38500 and BSU38510 and BSU38520 and BSU38530 | 3E7W;3E7X;&&4BPF;4BPG;4BPH;& | https://doi.org/10.1074/jbc.M800557200 https://doi.org/10.1016/j.febslet.2015.07.008 |
| BSU32490 or BSU32470 or BSU32510 or BSU32480 or BSU32500 |  |  |
| BSU32500 or BSU32470 or BSU32490 or BSU32510 or BSU32480 |  |  |
| BSU32820 or BSU24150 or BSU37170 or BSU18260 or BSU04520 | &5LNX;&& |  |
| BSU24970 and BSU24960 and BSU24950 and BSU24980 and BSU24990 |  |  |
| BSU25900 and BSU17410 and BSU01530 and BSU25710 and BSU35620 | &1X60;&& |  |
| BSU30450 and BSU30440 and BSU30430 and BSU30420 and BSU30410 |  |  |
| BSU30450 and BSU30440 and BSU30430 and BSU30420 and BSU30410 |  |  |
| BSU39670 and BSU39750 and BSU39740 and BSU39730 and BSU39720 |  |  |
| BSU31610 or BSU31660 or BSU31600 or BSU31620 or BSU31630 or BSU31650 |  |  |
| BSU35609 and BSU35600 and BSU35590 and BSU35570 and BSU35550 and BSU35540 |  |  |
| BSU35750 and BSU35760 and BSU35720 and BSU35710 and BSU35700 and BSU35530 | &&&&7DD0; |  |
| BSU35750 and BSU35760 and BSU35730 and BSU35720 and BSU35710 and BSU35700 and BSU35530 | &&&&&7DD0; |  |
| BSU36880 and BSU36870 and BSU36860 and BSU36850 and BSU36840 and BSU36830 and BSU36820 and BSU36810 and BSU36800 |  |  |
| BSU38500 and BSU38510 and BSU38520 and BSU38530 and BSU35750 and BSU35760 and BSU35720 and BSU35710 and BSU35700 and BSU35530 | 3E7W;3E7X;&&4BPF;4BPG;4BPH;&&&&&&7DD0; | https://doi.org/10.1074/jbc.M800557200 https://doi.org/10.1016/j.febslet.2015.07.008 https://doi.org/10.1016/j.bbrc.2020.12.028 |
| BSU31600 or BSU09850 or BSU31610 or BSU31620 or BSU31630 or BSU31640 or BSU31650 or BSU31660 or BSU09680 or BSU33420 or BSU11640 |  |  |
| BSU26690 or ( BSU26710 and BSU26700 ) or BSU29600 |  |  |
| BSU26690 or ( BSU26710 and BSU26700 ) or BSU29600 |  |  |
| ( BSU26710 and BSU26700 ) or BSU29600 or BSU26690 |  |  |
| ( BSU14580 and BSU14590 ) and BSU14600 and BSU14610 |  |  |
| ( BSU11230 and BSU11240 ) or ( BSU15510 and BSU15520 ) |  |  |
| ( BSU30710 and BSU30720 ) or ( BSU38760 and BSU38750 ) |  |  |
| ( BSU29200 and BSU29210 ) and BSU24350 and BSU24340 and BSU22440 |  |  |
| ( BSU31090 and BSU31100 ) or ( BSU14510 and BSU13500 ) or BSU26640 | 1LSU;2HMS;2HMT;2HMU;2HMV;2HMW;4J7C;4J90;4J91;5BUT;6S2J;6S5B;6S5C;6S5D;6S5E;6S5G;6S5N;6S5O;6S7R;8K16;8K1K;8K1S;8K1T;8K1U;8XMH;8XMI;&4J7C;5BUT;8K1S;8K1T;8K1U;8POO;8XMH;8XMI;&6I8V;&&&&&& | https://doi.org/10.1016/s0092-8674(02)00768-7 https://doi.org/10.1016/j.cell.2006.08.028 https://doi.org/10.1038/nature12055 https://doi.org/10.1371/journal.pbio.1002356 https://doi.org/10.7554/eLife.50661 https://doi.org/10.1038/s41467-024-48057-y https://doi.org/10.1073/pnas.2318666121 https://doi.org/10.1016/j.jsb.2019.02.002 |
| ( BSU03320 and BSU03310 ) or ( BSU37280 and BSU37270 and BSU37260 ) |  |  |
| ( BSU13900 and BSU13910 and BSU38050 ) or ( BSU13900 and BSU13910 and BSU01680 ) | 1JEM;1KKL;1KKM;1SPH;2FEP;2HID;2HPR;3OQM;3OQN;3OQO;&&&1JEM;1KKL;1KKM;1SPH;2FEP;2HID;2HPR;3OQM;3OQN;3OQO;&&3PPN;3PPO;3PPP;3PPQ;3PPR;&&& | [https://doi.org/10.1002/pro.5560061006 https://doi.org/10.1073/pnas.192368699 https://doi.org/10.1016/s0969-2126(94)00122-7 https://doi.org/10.1002/prot.21001 https://doi.org/10.1093/nar/gkq1177 https://doi.org/10.1042/BJ20102097](https://doi.org/10.1002/pro.5560061006) |
| ( BSU36710 and BSU12160 ) or ( BSU36710 and BSU27220 ) or ( BSU36710 and BSU18570 ) |  |  |
| ( BSU02980 and BSU02990 and BSU03000 ) or ( BSU33830 and BSU33820 and BSU33810 and BSU33800 ) | &&2B4L;2B4M;3CHG;5NXX;&&&3PPN;3PPO;3PPP;3PPQ;3PPR;&&& | https://doi.org/10.1016/j.jmb.2005.12.085 https://doi.org/10.1128/JB.00346-08 https://doi.org/10.1111/1462-2920.13999 <https://doi.org/10.1042/BJ20102097> |
| ( BSU02980 and BSU02990 and BSU03000 ) or ( BSU33830 and BSU33820 and BSU33810 and BSU33800 ) | &&2B4L;2B4M;3CHG;5NXX;&&&3PPN;3PPO;3PPP;3PPQ;3PPR;&&& | https://doi.org/10.1016/j.jmb.2005.12.085 https://doi.org/10.1128/JB.00346-08 https://doi.org/10.1111/1462-2920.13999 <https://doi.org/10.1042/BJ20102097> |
| ( BSU03590 and BSU03600 and BSU03610 ) or ( BSU27440 and BSU27450 and BSU27460 and BSU27430 ) |  |  |
| ( BSU29690 and BSU29700 and BSU29710 ) or ( BSU08060 and BSU08070 and BSU08080 and BSU08090 ) |  |  |
| ( BSU33830 and BSU33820 and BSU33810 and BSU33800 ) or ( BSU02980 and BSU02990 and BSU03000 ) | &&3PPN;3PPO;3PPP;3PPQ;3PPR;&&&&2B4L;2B4M;3CHG;5NXX;&& | https://doi.org/10.1016/j.jmb.2005.12.085 https://doi.org/10.1128/JB.00346-08 https://doi.org/10.1111/1462-2920.13999 <https://doi.org/10.1042/BJ20102097> |
| ( BSU29380 and BSU29370 and BSU29360 and BSU29350 and BSU29340 ) or ( BSU32730 and BSU32740 and BSU32750 ) | &2O1M;&&&&4GOT;&&& |  |
| ( BSU32730 and BSU32740 and BSU32750 ) or ( BSU29380 and BSU29370 and BSU29360 and BSU29350 and BSU29340 ) | 4GOT;&&&&2O1M;&&&& |  |
| ( BSU32730 and BSU32740 and BSU32750 ) or ( BSU29380 and BSU29370 and BSU29360 and BSU29350 and BSU29340 ) | 4GOT;&&&&2O1M;&&&& |  |
| ( BSU32730 and BSU32740 and BSU32750 ) or ( BSU29380 and BSU29370 and BSU29360 and BSU29350 and BSU29340 ) | 4GOT;&&&&2O1M;&&&& |  |
| ( BSU33730 and BSU33720 and BSU33710 and BSU33700 ) or ( BSU33830 and BSU33820 and BSU33810 and BSU33800 ) | &&3R6U;5NXY;6EYG;6EYH;6EYL;6EYQ;&&&&3PPN;3PPO;3PPP;3PPQ;3PPR;&& | <https://doi.org/10.1016/j.jmb.2011.05.037> https://doi.org/10.1111/1462-2920.13999 <https://doi.org/10.1042/BJ20102097> |
| ( BSU27070 and BSU27060 and BSU27050 and BSU27040 and BSU13900 and BSU13910 ) or ( BSU14400 and BSU13900 and BSU13910 ) | &1BLE;&&&1JEM;1KKL;1KKM;1SPH;2FEP;2HID;2HPR;3OQM;3OQN;3OQO;&&2R4Q;&1JEM;1KKL;1KKM;1SPH;2FEP;2HID;2HPR;3OQM;3OQN;3OQO;& | [https://doi.org/10.1006/jmbi.1997.1544  https://doi.org/10.1002/pro.5560061006 https://doi.org/10.1073/pnas.192368699 https://doi.org/10.1016/s0969-2126(94)00122-7 https://doi.org/10.1002/prot.21001 https://doi.org/10.1093/nar/gkq1177](https://doi.org/10.1006/jmbi.1997.1544) |
| ( BSU34610 and BSU34600 and BSU34590 ) or ( BSU34160 and BSU34150 and BSU34140 ) or ( BSU30290 and BSU30280 and BSU30270 ) | &&&&&&&&4R6H; |  |
| ( BSU15900 and BSU11340 and BSU15910 and BSU11720 and BSU04030 ) or ( BSU15900 and BSU11340 and BSU15910 and BSU10170 and BSU08650 and BSU36370 ) | &4LS5;4LS6;4LS7;4LS8;&&3OIF;3OIG;&&&4LS5;4LS6;4LS7;4LS8;&&8VDB;&3OIC;3OID; | [https://doi.org/10.1111/febs.12785 https://doi.org/10.1016/j.jmb.2010.12.003 https://doi.org/10.1016/j.jsb.2024.108065](https://doi.org/10.1111/febs.12785) <https://doi.org/10.1016/j.jmb.2010.12.003> |
| ( BSU15900 and BSU11340 and BSU15910 and BSU11720 and BSU04030 ) or ( BSU15900 and BSU11340 and BSU15910 and BSU10170 and BSU08650 and BSU36370 ) | &4LS5;4LS6;4LS7;4LS8;&&3OIF;3OIG;&&&4LS5;4LS6;4LS7;4LS8;&&8VDB;&3OIC;3OID; | [https://doi.org/10.1111/febs.12785 https://doi.org/10.1016/j.jmb.2010.12.003 https://doi.org/10.1016/j.jsb.2024.108065](https://doi.org/10.1111/febs.12785) <https://doi.org/10.1016/j.jmb.2010.12.003> |
| ( BSU15900 and BSU11340 and BSU15910 and BSU10170 and BSU08650 and BSU36370 ) or ( BSU15900 and BSU11340 and BSU15910 and BSU11330 and BSU11720 and BSU04030 ) | &4LS5;4LS6;4LS7;4LS8;&&8VDB;&3OIC;3OID;&&&4LS5;4LS6;4LS7;4LS8;&&8VD9;8VDA;&3OIF;3OIG; | [https://doi.org/10.1111/febs.12785 https://doi.org/10.1016/j.jmb.2010.12.003 https://doi.org/10.1016/j.jsb.2024.108065](https://doi.org/10.1111/febs.12785) <https://doi.org/10.1016/j.jmb.2010.12.003> |
| ( BSU15900 and BSU11340 and BSU15910 and BSU10170 and BSU08650 and BSU36370 ) or ( BSU15900 and BSU11340 and BSU15910 and BSU11330 and BSU11720 and BSU04030 ) | &4LS5;4LS6;4LS7;4LS8;&&8VDB;&3OIC;3OID;&&&4LS5;4LS6;4LS7;4LS8;&&8VD9;8VDA;&3OIF;3OIG; | [https://doi.org/10.1111/febs.12785 https://doi.org/10.1016/j.jmb.2010.12.003 https://doi.org/10.1016/j.jsb.2024.108065](https://doi.org/10.1111/febs.12785) <https://doi.org/10.1016/j.jmb.2010.12.003> |
| ( BSU15900 and BSU11340 and BSU15910 and BSU10170 and BSU08650 and BSU36370 ) or ( BSU15900 and BSU11340 and BSU15910 and BSU11330 and BSU11720 and BSU04030 ) | &4LS5;4LS6;4LS7;4LS8;&&8VDB;&3OIC;3OID;&&&4LS5;4LS6;4LS7;4LS8;&&8VD9;8VDA;&3OIF;3OIG; | [https://doi.org/10.1111/febs.12785 https://doi.org/10.1016/j.jmb.2010.12.003 https://doi.org/10.1016/j.jsb.2024.108065](https://doi.org/10.1111/febs.12785) <https://doi.org/10.1016/j.jmb.2010.12.003> |
| ( BSU15900 and BSU11340 and BSU15910 and BSU10170 and BSU08650 and BSU36370 ) or ( BSU15900 and BSU11340 and BSU15910 and BSU11330 and BSU11720 and BSU04030 ) | &4LS5;4LS6;4LS7;4LS8;&&8VDB;&3OIC;3OID;&&&4LS5;4LS6;4LS7;4LS8;&&8VD9;8VDA;&3OIF;3OIG; | [https://doi.org/10.1111/febs.12785 https://doi.org/10.1016/j.jmb.2010.12.003 https://doi.org/10.1016/j.jsb.2024.108065](https://doi.org/10.1111/febs.12785) <https://doi.org/10.1016/j.jmb.2010.12.003> |
| ( BSU15900 and BSU11340 and BSU15910 and BSU11330 and BSU11720 and BSU04030 ) or ( BSU15900 and BSU11340 and BSU15910 and BSU10170 and BSU08650 and BSU36370 ) | &4LS5;4LS6;4LS7;4LS8;&&8VD9;8VDA;&3OIF;3OIG;&&&4LS5;4LS6;4LS7;4LS8;&&8VDB;&3OIC;3OID; | [https://doi.org/10.1111/febs.12785 https://doi.org/10.1016/j.jmb.2010.12.003 https://doi.org/10.1016/j.jsb.2024.108065](https://doi.org/10.1111/febs.12785) <https://doi.org/10.1016/j.jmb.2010.12.003> |
| ( BSU15900 and BSU11340 and BSU15910 and BSU11330 and BSU11720 and BSU04030 ) or ( BSU15900 and BSU11340 and BSU15910 and BSU10170 and BSU08650 and BSU36370 ) | &4LS5;4LS6;4LS7;4LS8;&&8VD9;8VDA;&3OIF;3OIG;&&&4LS5;4LS6;4LS7;4LS8;&&8VDB;&3OIC;3OID; | [https://doi.org/10.1111/febs.12785 https://doi.org/10.1016/j.jmb.2010.12.003 https://doi.org/10.1016/j.jsb.2024.108065](https://doi.org/10.1111/febs.12785) <https://doi.org/10.1016/j.jmb.2010.12.003> |
| ( BSU15900 and BSU11340 and BSU15910 and BSU11330 and BSU11720 and BSU04030 ) or ( BSU15900 and BSU11340 and BSU15910 and BSU10170 and BSU08650 and BSU36370 ) | &4LS5;4LS6;4LS7;4LS8;&&8VD9;8VDA;&3OIF;3OIG;&&&4LS5;4LS6;4LS7;4LS8;&&8VDB;&3OIC;3OID; | [https://doi.org/10.1111/febs.12785 https://doi.org/10.1016/j.jmb.2010.12.003 https://doi.org/10.1016/j.jsb.2024.108065](https://doi.org/10.1111/febs.12785) <https://doi.org/10.1016/j.jmb.2010.12.003> |
| ( BSU15900 and BSU11340 and BSU15910 and BSU11330 and BSU11720 and BSU04030 ) or ( BSU15900 and BSU11340 and BSU15910 and BSU10170 and BSU08650 and BSU36370 ) | &4LS5;4LS6;4LS7;4LS8;&&8VD9;8VDA;&3OIF;3OIG;&&&4LS5;4LS6;4LS7;4LS8;&&8VDB;&3OIC;3OID; | [https://doi.org/10.1111/febs.12785 https://doi.org/10.1016/j.jmb.2010.12.003 https://doi.org/10.1016/j.jsb.2024.108065](https://doi.org/10.1111/febs.12785) <https://doi.org/10.1016/j.jmb.2010.12.003> |
| ( BSU15900 and BSU11340 and BSU15910 and BSU11330 and BSU11720 and BSU04030 ) or ( BSU15900 and BSU11340 and BSU15910 and BSU10170 and BSU08650 and BSU36370 ) | &4LS5;4LS6;4LS7;4LS8;&&8VD9;8VDA;&3OIF;3OIG;&&&4LS5;4LS6;4LS7;4LS8;&&8VDB;&3OIC;3OID; | [https://doi.org/10.1111/febs.12785 https://doi.org/10.1016/j.jmb.2010.12.003 https://doi.org/10.1016/j.jsb.2024.108065](https://doi.org/10.1111/febs.12785) <https://doi.org/10.1016/j.jmb.2010.12.003> |
| ( BSU15900 and BSU11340 and BSU15910 and BSU11330 and BSU11720 and BSU04030 ) or ( BSU15900 and BSU11340 and BSU15910 and BSU10170 and BSU08650 and BSU36370 ) | &4LS5;4LS6;4LS7;4LS8;&&8VD9;8VDA;&3OIF;3OIG;&&&4LS5;4LS6;4LS7;4LS8;&&8VDB;&3OIC;3OID; | [https://doi.org/10.1111/febs.12785 https://doi.org/10.1016/j.jmb.2010.12.003 https://doi.org/10.1016/j.jsb.2024.108065](https://doi.org/10.1111/febs.12785) <https://doi.org/10.1016/j.jmb.2010.12.003> |
| ( BSU15900 and BSU11340 and BSU15910 and BSU11330 and BSU11720 and BSU04030 ) or ( BSU15900 and BSU11340 and BSU15910 and BSU10170 and BSU08650 and BSU36370 ) | &4LS5;4LS6;4LS7;4LS8;&&8VD9;8VDA;&3OIF;3OIG;&&&4LS5;4LS6;4LS7;4LS8;&&8VDB;&3OIC;3OID; | [https://doi.org/10.1111/febs.12785 https://doi.org/10.1016/j.jmb.2010.12.003 https://doi.org/10.1016/j.jsb.2024.108065](https://doi.org/10.1111/febs.12785) <https://doi.org/10.1016/j.jmb.2010.12.003> |
| ( BSU15900 and BSU11340 and BSU15910 and BSU11330 and BSU11720 and BSU04030 ) or ( BSU15900 and BSU11340 and BSU15910 and BSU10170 and BSU08650 and BSU36370 ) | &4LS5;4LS6;4LS7;4LS8;&&8VD9;8VDA;&3OIF;3OIG;&&&4LS5;4LS6;4LS7;4LS8;&&8VDB;&3OIC;3OID; | [https://doi.org/10.1111/febs.12785 https://doi.org/10.1016/j.jmb.2010.12.003 https://doi.org/10.1016/j.jsb.2024.108065](https://doi.org/10.1111/febs.12785) <https://doi.org/10.1016/j.jmb.2010.12.003> |
| ( BSU07520 and BSU07510 and BSU07500 and BSU07490 ) or ( BSU33310 and BSU33290 and BSU33320 and BSU33300 ) or ( BSU03800 and BSU03810 and BSU03820 and BSU03830 ) | &&&&&&3G9Q;3HXP;&&&&&3GFV; |  |
| ( BSU13900 and BSU13910 and BSU38390 ) or ( BSU13900 and BSU13910 and BSU05810 and BSU05820 and BSU05830 ) or ( BSU38570 and BSU38590 and BSU38580 and BSU13900 and BSU13910 ) | 1JEM;1KKL;1KKM;1SPH;2FEP;2HID;2HPR;3OQM;3OQN;3OQO;&&&1JEM;1KKL;1KKM;1SPH;2FEP;2HID;2HPR;3OQM;3OQN;3OQO;&&&&&&&&1JEM;1KKL;1KKM;1SPH;2FEP;2HID;2HPR;3OQM;3OQN;3OQO; | [https://doi.org/10.1002/pro.5560061006 https://doi.org/10.1073/pnas.192368699 https://doi.org/10.1016/s0969-2126(94)00122-7 https://doi.org/10.1002/prot.21001 https://doi.org/10.1093/nar/gkq1177 https://doi.org/10.1042/BJ20102097](https://doi.org/10.1002/pro.5560061006) |
| ( BSU28310 and BSU28300 ) or BSU36010 |  |  |
| BSU36010 or ( BSU28310 and BSU28300 ) |  |  |
| BSU29020 | 8WWZ; | <https://doi.org/10.34184/kssb.2023.11.4.59> |
| BSU03930 | 8W0N; | <https://doi.org/10.1021/acscatal.3c05615> |
| BSU10170 | 8VDB; | <https://doi.org/10.1016/j.jsb.2024.108065> |
| BSU11330 | 8VD9;8VDA; | <https://doi.org/10.1016/j.jsb.2024.108065> |
| BSU21810 | 8UVZ; |  |
| BSU12960 | 8AY0;8AZB; | <https://doi.org/10.1099/mic.0.001274> |
| BSU00740 | 7PI1; |  |
| BSU01770 | 7OJR;7OJS;7OLH;7OML; |  |
| BSU00090 | 7OJ1;7OJ2; |  |
| BSU18440 | 7MFT; |  |
| BSU18450 | 7MFT; |  |
| BSU16530 | 7JLI;7JLJ;7JLM;7JLR; |  |
| BSU35700 | 7DD0; |  |
| BSU00180 | 7CPH; |  |
| BSU18800 | 6W2Z; |  |
| BSU38170 | 6KOB;6KOC;6KOE; |  |
| BSU38150 | 6KOB;6KOC;6KOE; |  |
| BSU38160 | 6KOB;6KOC;6KOE; |  |
| BSU38140 | 6KOB;6KOC;6KOE; |  |
| BSU38260 | 6KMM;6KMN;7DLK;7E5Q;7PKX;7PL0; |  |
| BSU06240 | 6IE0; |  |
| BSU14510 | 6I8V; |  |
| BSU27600 | 6HTQ;6YXA;8ACU; |  |
| BSU22480 | 6F2C; |  |
| BSU40320 | 6DKT;6NFP; |  |
| BSU17380 | 6CGL;6CGM;6CGN;6MT9;6MV9;6MVE;6MW3;6MYX; |  |
| BSU32450 | 6A4M; |  |
| BSU14870 | 6A2J;6IED; |  |
| BSU15620 | 5ZT7;5ZT8;5ZT9;5ZTA; |  |
| BSU36000 | 5XNE; |  |
| BSU03270 | 5XHU; |  |
| BSU02130 | 5T91;5T9B;5T9C; |  |
| BSU24130 | 5MUX; |  |
| BSU24170 | 5LP7; |  |
| BSU24150 | 5LNX; |  |
| BSU00910 | 5IWX;5IWY; |  |
| BSU29750 | 5GMU;5GO2; |  |
| BSU30240 | 5FLG;5FLL;5FM0;5G1F; |  |
| BSU39400 | 5EP8;5OLN; |  |
| BSU02090 | 5E2F;6W5E;6W5F; |  |
| BSU00900 | 5DDT;5DDV;5HS2; |  |
| BSU30790 | 5BUQ;5BUR;5BUS;5GTD;5X8F;5X8G; |  |
| BSU17150 | 4YXQ;4YXT;4YXV; |  |
| BSU06970 | 4R6K;5Z6B;5Z6C; |  |
| BSU30270 | 4R6H; |  |
| BSU34560 | 4M56;4M8U;4MAZ;4MB1;5WCZ;7LV6; |  |
| BSU11340 | 4LS5;4LS6;4LS7;4LS8; |  |
| BSU17460 | 4LNF;4LNI;4LNK;4LNN;4LNO;4S0R; |  |
| BSU07920 | 4KJR;4KJS; |  |
| BSU31100 | 4J7C;5BUT;8K1S;8K1T;8K1U;8POO;8XMH;8XMI; |  |
| BSU32730 | 4GOT; |  |
| BSU17390 | 4DR0; |  |
| BSU38830 | 4DNG; |  |
| BSU19630 | 4D8V;4D8X;4D8Y;4D98;4D9H;4DA0;4DA6;4DA7;4DA8;4DAB;4DAE;4DAN;4DAO;4DAR; |  |
| BSU38520 | 4BPF;4BPG;4BPH; |  |
| BSU29190 | 4A3S; | <https://doi.org/10.1016/j.jmb.2011.12.024> |
| BSU33900 | 4A3R;7XML; | <https://doi.org/10.1016/j.jmb.2011.12.024> |
| BSU04740 | 3W40;3W41;3W42;3W43;3W44;3W45; |  |
| BSU37710 | 3VMM;3WNZ;3WO0;3WO1; |  |
| BSU37110 | 3R8R; |  |
| BSU33710 | 3R6U;5NXY;6EYG;6EYH;6EYL;6EYQ; |  |
| BSU03050 | 3PQD;3PQE;3PQF; |  |
| BSU33810 | 3PPN;3PPO;3PPP;3PPQ;3PPR; |  |
| BSU11720 | 3OIF;3OIG; |  |
| BSU08650 | 3OIC;3OID; |  |
| BSU34550 | 3NAS; |  |
| BSU38110 | 3N2S; |  |
| BSU39700 | 3MZ0;3NT2;3NT4;3NT5;3NTO;3NTQ;3NTR;4L8V;4L9R; |  |
| BSU39380 | 3M1R; |  |
| BSU32110 | 3LZW;3LZX; |  |
| BSU37790 | 3K92; |  |
| BSU22960 | 3K8Z;7MFT; |  |
| BSU10140 | 3I6D; |  |
| BSU07840 | 3GVE; |  |
| BSU03830 | 3GFV; |  |
| BSU33320 | 3G9Q;3HXP; |  |
| BSU15110 | 3EGO; |  |
| BSU38500 | 3E7W;3E7X; |  |
| BSU30230 | 3DOD;3DRD;3DU4;6WNN; |  |
| BSU33400 | 3D3F;3F7J; |  |
| BSU38980 | 3CDK; |  |
| BSU38990 | 3CDK; |  |
| BSU28120 | 3BS8; |  |
| BSU29050 | 3B3D; |  |
| BSU13590 | 2ZVI; |  |
| BSU13550 | 2YRF;2YVK; |  |
| BSU06530 | 2XCL;2XD4; |  |
| BSU17660 | 2XCD;2XCE;4AOO;4AOZ;4APZ;4B0H; |  |
| BSU30820 | 2X7J; |  |
| BSU35010 | 2VHL; |  |
| BSU34920 | 2VD2; |  |
| BSU18410 | 2V36;3A75;3WHQ;3WHR;3WHS; |  |
| BSU16730 | 2RIR; |  |
| BSU25020 | 2RCV; |  |
| BSU14400 | 2R4Q; |  |
| BSU12010 | 2R48; |  |
| BSU01630 | 2PHZ;2WHY;2WI8;2XUZ;2XV1; |  |
| BSU39540 | 2OQC; |  |
| BSU13560 | 2OLC;2PU8;2PUI;2PUL;2PUN;2PUP; |  |
| BSU29370 | 2O1M; |  |
| BSU02850 | 2O1E; |  |
| BSU00110 | 2NV1;2NV2; |  |
| BSU21680 | 2KZN;3E0O; |  |
| BSU10120 | 2INF; |  |
| BSU38020 | 2I5B; |  |
| BSU32460 | 2H0E;2H0F;2H0J; |  |
| BSU39360 | 2FKN; |  |
| BSU13600 | 2FEA; |  |
| BSU09440 | 2C6X; |  |
| BSU35020 | 2BKV;2BKX; |  |
| BSU39370 | 2BB0;2G3F; |  |
| BSU20020 | 2BAZ;2XX6;2XY3;2Y1T;4AO5; |  |
| BSU03000 | 2B4L;2B4M;3CHG;5NXX; |  |
| BSU23280 | 2B3Z;2D5N;3EX8;4G3M; |  |
| BSU15490 | 2AT2;3R7D;3R7F;3R7L; |  |
| BSU28390 | 1ZUW; |  |
| BSU03860 | 1ZCH; |  |
| BSU18230 | 1YDO; |  |
| BSU39980 | 1Y3T;2H0V;8HFB; |  |
| BSU22070 | 1Y0B;2FXV;6W1I; |  |
| BSU05860 | 1XC3;3LM9;3OHR; | <https://doi.org/10.1016/j.jmb.2010.12.021> |
| BSU17410 | 1X60; |  |
| BSU08180 | 1U8X; |  |
| BSU11650 | 1TO9;1TYH;1YAF;1YAK;2QCX; |  |
| BSU13170 | 1TIY;1WKQ; |  |
| BSU37660 | 1TD9;1XCO; |  |
| BSU39760 | 1T90; |  |
| BSU06460 | 1T4A;1TWJ; |  |
| BSU23250 | 1RVV;1ZIS; |  |
| BSU17370 | 1RLJ; |  |
| BSU00120 | 1R9G;2NV0;2NV2; |  |
| BSU35790 | 1QWR; |  |
| BSU09530 | 1PZ1; |  |
| BSU39780 | 1PYF;1PZ0; |  |
| BSU01370 | 1P3J;2EU8;2OO7;2ORI;2OSB;2P3S;2QAJ;3DKV;3DL0;4MKF;4MKG;4MKH;4QBF;4QBG;4TYP;4TYQ;5X6I; |  |
| BSU22870 | 1P0K;1P0N; |  |
| BSU35930 | 1OGC;1OGD;1OGE;1OGF; |  |
| BSU35660 | 1O6C;4FKZ; |  |
| BSU15020 | 1O6B; |  |
| BSU11670 | 1NG3;1NG4;1RYI;3IF9; |  |
| BSU02430 | 1MKI;2OSU;3AGF;3BRM; |  |
| BSU31980 | 1MD9;1MDB;1MDF; |  |
| BSU31090 | 1LSU;2HMS;2HMT;2HMU;2HMV;2HMW;4J7C;4J90;4J91;5BUT;6S2J;6S5B;6S5C;6S5D;6S5E;6S5G;6S5N;6S5O;6S7R;8K16;8K1K;8K1S;8K1T;8K1U;8XMH;8XMI; |  |
| BSU25640 | 1KAM;1KAQ; |  |
| BSU40550 | 1K23;1WPM;1WPN;2HAW;2IW4; |  |
| BSU33510 | 1K0V;1P8G;2QIF;3I9Z; |  |
| BSU33500 | 1JWW;1KQK;1OPZ;1OQ3;1OQ6;1P6T;2RML;2VOY; |  |
| BSU25300 | 1JTK;1UWZ;1UX0;1UX1; |  |
| BSU13900 | 1JEM;1KKL;1KKM;1SPH;2FEP;2HID;2HPR;3OQM;3OQN;3OQO; |  |
| BSU33240 | 1J58;1L3J;1UW8;2UY8;2UY9;2UYA;2UYB;2V09;3S0M;4MET;5HI0;5VG3;6TZP;6UFI; |  |
| BSU37500 | 1IY9; |  |
| BSU29130 | 1HQS; |  |
| BSU12920 | 1HI9; |  |
| BSU38290 | 1G4E;1G4P;1G4S;1G4T;1G67;1G69;1G6C;2TPS;3O15;3O16; |  |
| BSU04620 | 1F7L;1F7T;1F80; |  |
| BSU06440 | 1F1O; |  |
| BSU03130 | 1EE1;1FYD;1IFX;1IH8;1KQP;1NSY;2NSY; |  |
| BSU00510 | 1DKR;1DKU;1IBS; |  |
| BSU15550 | 1DBT; |  |
| BSU35740 | 1COZ;1N1D; |  |
| BSU22690 | 1COM;1DBF;1FNJ;1FNK;2CHS;2CHT;3ZO8;3ZOP;3ZP4;3ZP7; |  |
| BSU38300 | 1C3Q;1EKK;1EKQ;1ESJ;1ESQ; |  |
| BSU27060 | 1BLE; | <https://doi.org/10.1006/jmbi.1997.1544> |
| BSU03040 | 1BAG;1UA7; |  |
| BSU17680 | 1B02;1BKO;1BKP;1BSF;1BSP; |  |
| BSU13890 | 1AX3;1GPR; |  |
| BSU06490 | 1AO0;1GPH; |  |
| BSU10130 | 1AK1;1C1H;1C9E;1DOZ;1LD3;1N0I;2AC2;2AC4;2H1V;2H1W;2HK6;2Q2N;2Q2O;2Q3J;3GOQ;3M4Z; |  |
| BSU34450 | 1OYG; 1PT2; 2VDT; 3BYJ; 3BYK; 3BYL; 3BYN; 6PWQ; 6VHQ; | https://doi.org/10.1038/nsb974 https://doi.org/10.1093/protein/gzn036 https://doi.org/10.1186/1472-6807-8-16 https://doi.org/10.1016/j.ijbiomac.2020.06.114 <https://doi.org/10.1074/jbc.RA120.015853> |
